# Biomolecular condensation orchestrates clathrin-mediated endocytosis in plants

**DOI:** 10.1101/2022.03.17.484738

**Authors:** Jonathan Michael Dragwidge, Yanning Wang, Lysiane Brocard, Andreas De Meyer, Roman Hudeček, Dominique Eeckhout, Peter Grones, Matthieu Buridan, Clément Chambaud, Přemysl Pejchar, Martin Potocký, Joanna Winkler, Michael Vandorpe, Nelson Serre, Matyáš Fendrych, Amelie Bernard, Geert De Jaeger, Roman Pleskot, Xiaofeng Fang, Daniël Van Damme

**Affiliations:** Department of Plant Biotechnology and Bioinformatics, Ghent University, Technologiepark 71, 9052 Ghent, Belgium; VIB Center for Plant Systems Biology, Technologiepark 71, 9052 Ghent, Belgium; Center for Plant Biology, School of Life Sciences, Tsinghua University, Beijing, China; Univ. Bordeaux, CNRS, INSERM, Bordeaux Imaging Center, BIC, UAR 3420, US 4, F-33000 Bordeaux, France; Institute of Experimental Botany of the Czech Academy of Sciences, Rozvojová 263, 16502 Prague 6, Czech Republic; Laboratoire de Biogenèse Membranaire, UMR 5200, CNRS, Univ. Bordeaux, F-33140 Villenave d’Ornon; Department of Experimental Plant Biology, Faculty of Sciences, Charles University, Prague, Czech Republic

**Keywords:** Phase separation, Biomolecular condensate, Endocytosis, Membrane trafficking, Arabidopsis thaliana, TPLATE complex, clathrin

## Abstract

Clathrin-mediated endocytosis (CME) is an essential cellular internalisation pathway involving the dynamic assembly of clathrin and accessory proteins to form membrane-bound vesicles. In plants, the evolutionarily ancient TSET/TPLATE complex (TPC) plays an essential, but not well-defined role in CME. Here, we show that two highly disordered TPC subunits, AtEH1 and AtEH2 function as scaffolds to drive biomolecular condensation of the complex. These condensates specifically nucleate on the plasma membrane through interactions with anionic phospholipids, and facilitate the dynamic recruitment and assembly of clathrin, early-, and late-stage endocytic accessory proteins. Importantly, clathrin forms ordered assemblies within the condensate environment. Biomolecular condensation therefore acts to promote dynamic protein assemblies throughout clathrin-mediated endocytosis. Furthermore, the disordered region sequence properties of AtEH1 regulate the material properties of the endocytic condensates *in vivo* and alteration of these material properties influences endocytosis dynamics, and consequently plant adaptive growth.

**Highlights:**

- AtEH subunits are endocytic scaffolds which drive condensation of the TPC
- AtEH1 condensates nucleate on the plasma membrane via lipid interactions
- Condensation of AtEH1/TPC facilitates clathrin re-arrangement and assembly
- AtEH1 IDR1 composition controls condensate properties to regulate endocytosis

## Introduction

Clathrin-mediated endocytosis is an essential cellular internalisation pathway necessary for many cellular processes, including cell signalling, immune responses, and nutrient uptake (Kaksonen and Roux, 2018; McMahon and Boucrot, 2011; Paez Valencia et al., 2016). Endocytosis is initiated through a network of transient interactions between cargo, adaptor proteins, membrane lipids, and the coat protein clathrin on the plasma membrane. Following initiation, clathrin polymerises into a lattice which bends as the membrane invaginates, leading to dynamin-dependent scission and release of the formed clathrin-coated vesicle (CCV). The assembly of these clathrin lattices is highly dynamic, and depends on a network of early accessory proteins which interact with the adaptor protein complex AP-2 to stabilise clathrin-coated pits (Chen and Schmid, 2020; Sochacki and Taraska, 2019). Core endocytic machinery, including clathrin and AP-2 was present in the last eukaryotic common ancestor (Dacks and Robinson, 2017; Rout and Field, 2017), suggesting that the underlying principles that facilitate clathrin-coat formation during endocytosis are largely conserved between eukaryotes.

One key evolutionary difference is that the amoeba *Dictyostelium discoideum* and plants retained an ancient endocytic complex termed TSET, or TPLATE complex (TPC), which was lost from metazoan and fungal lineages (Gadeyne et al., 2014; Hirst et al., 2014). TPC is essential for plant life, as viable knock-out mutants have not been identified (Gadeyne et al., 2014; Van Damme et al., 2006; Wang et al., 2019). Recent studies have established the critical role of TPC during endocytosis by demonstrating its interaction with many essential endocytic components, including anionic phospholipids (Yperman et al., 2021a; Yperman et al., 2021b), the AP-2 complex (Gadeyne et al., 2014; Wang et al., 2023), clathrin (Van Damme et al., 2011), and ubiquitinated cargo (Grones et al., 2022). Furthermore, destabilisation of TPC leads to impaired endocytic internalisation, and disrupts membrane bending during clathrin-coated pit formation (Johnson et al., 2021; Wang et al., 2021). TPC is present throughout endocytosis progression, from the earliest detectable phase until vesicle scission and uncoating (Gadeyne et al., 2014; Narasimhan et al., 2020). Collectively, these studies support a multi-functional role for TPC during endocytosis.

While TPC shares structural homology with COPI and AP-complexes (Rout and Field, 2017; Yperman et al., 2021b), TPC subunits do not have direct orthologs to human and fungal proteins. However, TPC subunits contain common vesicle trafficking related domains and motifs which has provided insight into their molecular function (Zhang et al., 2015). Notably, the plant specific TPC subunits AtEH1/Pan1 (AtEH1) and AtEH2/Pan1 (AtEH2) share partial homology to the Eps15 homology (EH) domains proteins Eps15 and Intersectin in humans, and Ede1p and Pan1p in yeast. A conserved function in autophagy of AtEH proteins (Wang et al., 2019), and Ede1p and Pan1p in yeast (Liu et al., 2022; Wilfling et al., 2020) support the idea that they are functional homologs. We previously showed that AtEH1 co-purifies with AP-2 (Gadeyne et al., 2014; Wang et al., 2023), and AtEH1 interacts with anionic phospholipids and cargo (Yperman et al., 2021a). Interactomics and live cell imaging suggest TPC assembles as an octameric complex on the membrane during the initiation of endocytosis (Gadeyne et al., 2014; Wang et al., 2020), with the lipid binding proteins AtEH1, AtEH2, and the muniscin-like protein TML forming an interface with the plasma membrane (Yperman et al., 2021b). In comparison, human Eps15, intersectin and the muniscin FCHo mediate multivalent interactions between themselves, anionic phospholipids (Alaoui et al., 2022; Day et al., 2021; Henne et al., 2010), and AP-2 (Hollopeter et al., 2014; Ma et al., 2015; Partlow et al., 2022) to promote the initiation and growth of clathrin coated pits during endocytosis (Bhave et al., 2020; Cocucci et al., 2012; Lehmann et al., 2019). Thus, AtEH1, AtEH2 and TML may be plant endocytic initiation components.

Recently, membraneless organelles (or biomolecular condensates), which can be assembled through phase separation has emerged as a fundamental mechanism to compartmentalise cellular functions (Banani et al., 2017; Choi et al., 2020; Snead and Gladfelter, 2019; Zhao and Zhang, 2020). In animal and yeast, Eps15/FCHo and Ede1 have been shown to promote the formation of condensates during the early phase of endocytosis (Day et al., 2021; Kozak and Kaksonen, 2022). Here, we demonstrate that in plants, AtEH proteins form condensates on the plasma membrane to recruit cytosolic proteins throughout endocytosis. Moreover, using CLEM-ET we found that clathrin forms lattices within the liquid-like assemblies generated by AtEH1/TPC. Evolutionarily analysis revealed that the physical properties of TPC condensates are determined from the sequence chemistry of the intrinsically disordered region 1 (IDR1) of AtEH1. Changes to these properties alter endocytic kinetics *in vivo*. Our findings provide general insight on how collective interactions shape endocytosis in eukaryotes.

## Results

### AtEH1 and AtEH2 phase separate *in vivo* and *in vitro*

Yeast Ede1 and human Eps15 are key endocytic components containing EH domains and intrinsic disordered regions, and have been shown to phase separate (Day et al., 2021; Kozak and Kaksonen, 2022; Wilfling et al., 2020). Sequence analysis revealed that AtEH1 and AtEH2 are highly disordered compared to other TPC subunits (Figures S1A and S1B), containing three intrinsically disordered regions (IDRs) and prion-like regions (Figure 1A), common features of proteins which phase separate (Alberti et al., 2019). Because phase separation is concentration dependent, we tested whether AtEH1 and AtEH2 may form condensates when over-expressed in cells. We observed punctate cytosolic and membrane associated puncta after transient overexpression of AtEH proteins in tobacco (*Nicotiana benthamiana*), or stable overexpression in *Arabidopsis thaliana* (Figure 1B), consistent with previous studies (Wang et al., 2019). Similar compartments were observed by heterologous expression of AtEH proteins in fission yeast (*Schizosaccharomyces pombe*) (Figure 1B). We next asked whether these punctate assemblies have physical properties consistent with biomolecular condensates. At very high expression levels we observed the dynamic growth and shrinking of puncta consistent with Ostwald ripening (Figure 1C), as well as fusion of puncta (Figure 1D). Furthermore, fluorescence recovery after photobleaching (FRAP) experiments indicate that AtEH1-GFP molecules are rapidly exchanged between the puncta and the cytosolic pool of proteins (Figure 1E). Collectively, these properties are consistent with protein phase separation, and indicate that AtEH1 and AtEH2 have the capacity to form biomolecular condensates *in vivo*. Consequently, our results indicate that the previously identified co-localisation between AtEH1 and ATG8 in *N. benthamiana* (Wang et al., 2019) indicates partitioning of ATG8 with AtEH1 condensates, rather than induction of autophagosomes in that system.

**Figure 1.**
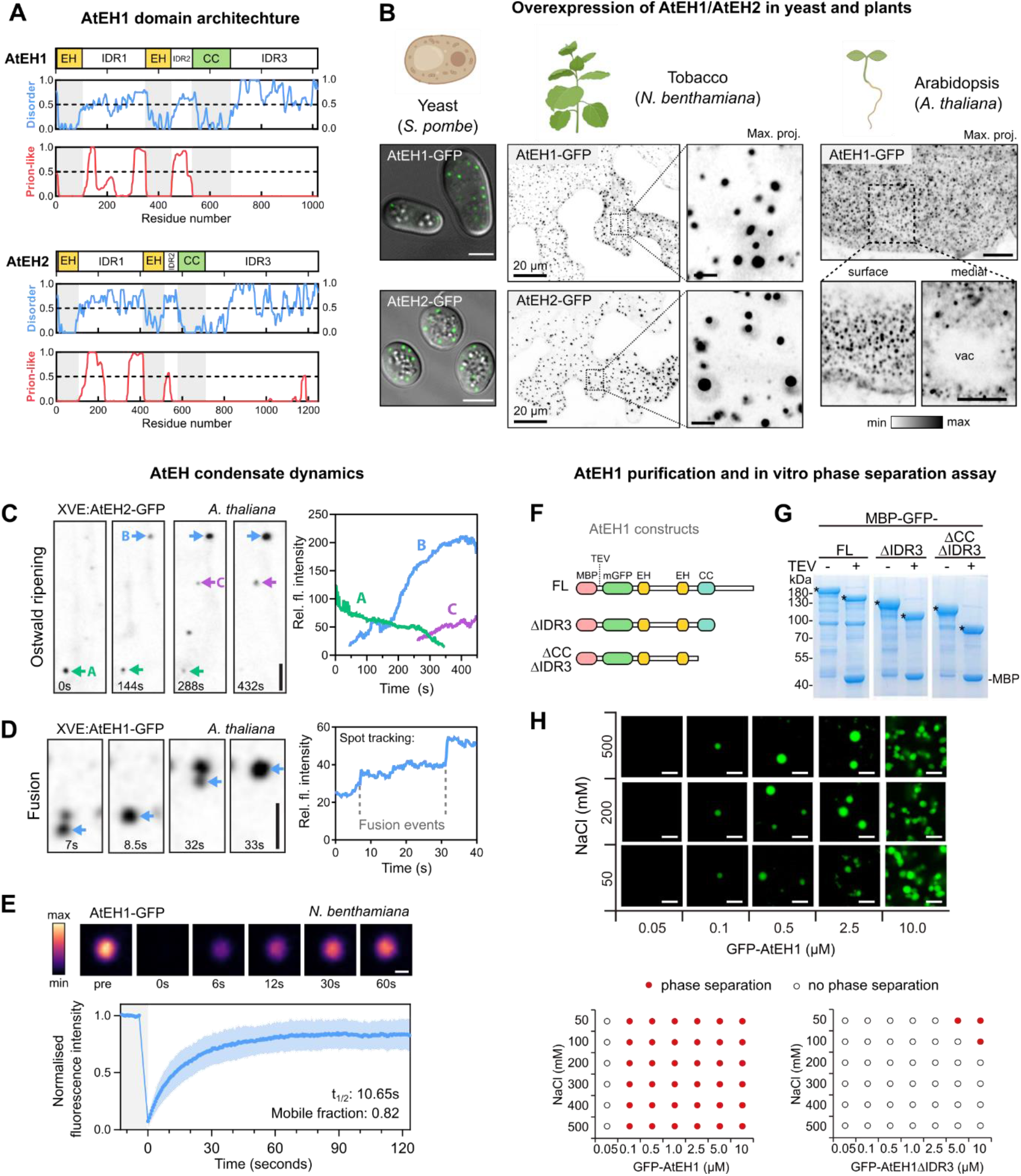
AtEH/Pan1 proteins phase separate *in vivo* and *in vitro*. (A) AtEH1 and AtEH2 domain architecture and prediction of disordered (MobiDB consensus) and prion-like (PLAAC) residues. Regions with values > 0.5 are considered disordered, or prion-like respectively. EH, Eps15 homology; CC, coiled-coil. (B) Airyscan images of AtEH1-GFP and AtEH2-GFP overexpressed in yeast (*S. pombe*), *N. benthamiana* epidermal cells (UBQ10:AtEH-mGFP), and in stable *A. thaliana* root epidermal cells (35S:AtEH1-GFP). (C-D) Time-lapse imaging of AtEH1 and AtEH2 under control of an estradiol inducible promoter (XVE) after 38 (C) or 22 (D) hours induction in *A. thaliana* root epidermal cells. Spot tracking and quantification of fluorescence intensity reveals puncta growth and shrinking through Ostwald ripening (C), and puncta fusion (D). (E) Fluorescence recovery after photobleaching (FRAP) of AtEH1-GFP condensates in *N. benthamiana* epidermal cells. Data is mean ± SD, n= 20 spots from 12 cells. (F-H) Schematic of AtEH1 constructs used for recombinant protein purification (F). Constructs were purified as Green Fluorescent protein (GFP)-fusion proteins using an N-terminally located Maltose Binding Protein (MBP) tag with a Tobacco Etch Virus (TEV) cleavage site. (G) Coomassie-stained SDS-PAGE gel of purified AtEH1 protein before and after TEV cleavage; * indicates AtEH1. (H) *in vitro* phase separation assay and phase diagram of recombinant GFP-AtEH1. GFP-AtEH1_ΔCCΔIDR3_ did not phase separate at the tested conditions. Scale bars = 5 μm (B, C, D), 1 μm (E), 2 μm (H), or as otherwise indicated. **See also Figure S1.**

Since AtEH1 and AtEH2 have a similar structure and ability to phase separate, we focused our work on AtEH1 due to its more central connection to TML and TPLATE in TPC compared to AtEH2 (Yperman et al., 2021b). We next examined whether AtEH1 can phase separate *in vitro*. We generated three AtEH1 constructs fused to MBP-TEV-GFP: full length (AtEH1_FL_), an IDR3 truncation (AtEH1_ΔIDR3_), and a coiled-coil (CC) and IDR3 truncation (AtEH1_ΔCCΔIDR3_) (Figure 1F). We expressed and purified these proteins recombinantly in *E. coli* (Figure 1G), and then performed *in vitro* phase separation assays under different salt and protein concentrations (Figure 1H). AtEH1_FL_ underwent phase separation at relatively low protein concentrations (≤ 0.1 µM), but showed relatively limited fluorescence recovery (Figure S1C). This may suggest that intra-molecular interactions promote phase separation but limit molecular re-arrangement in the absence of other TPC components. Additionally, phase separation was drastically reduced when the IDR3 was truncated (AtEH1_ΔIDR3_), and abolished when the coiled-coil domain was further truncated (AtEH1_ΔCCΔIDR3_) (Figure 1H), indicating that the coiled-coil and IDR3 are important domains promoting phase separation in the absence of other factors.

### Condensation of AtEH1 is controlled by multiple regions, including IDR1

We next asked whether we could determine the region(s) which promote condensation of AtEH1 *in vivo*. To test this, we systematically removed each intrinsically disordered region (IDR) and structured domain in AtEH1 and quantified the relative concentration of protein in the cytosol (light phase; C_L_), as a proxy for protein saturation concentration (C_sat_) (Figures 2A-B and S1A). Deletion of all domains resulted in significantly increased C_L_, implying that both IDRs and structured domains promote condensation *in vivo*. Notably, deletion of IDR1 resulted in abnormally formed condensates, suggesting that IDR1 may function as a regulatory region (Figures 2A).

**Figure 2.**
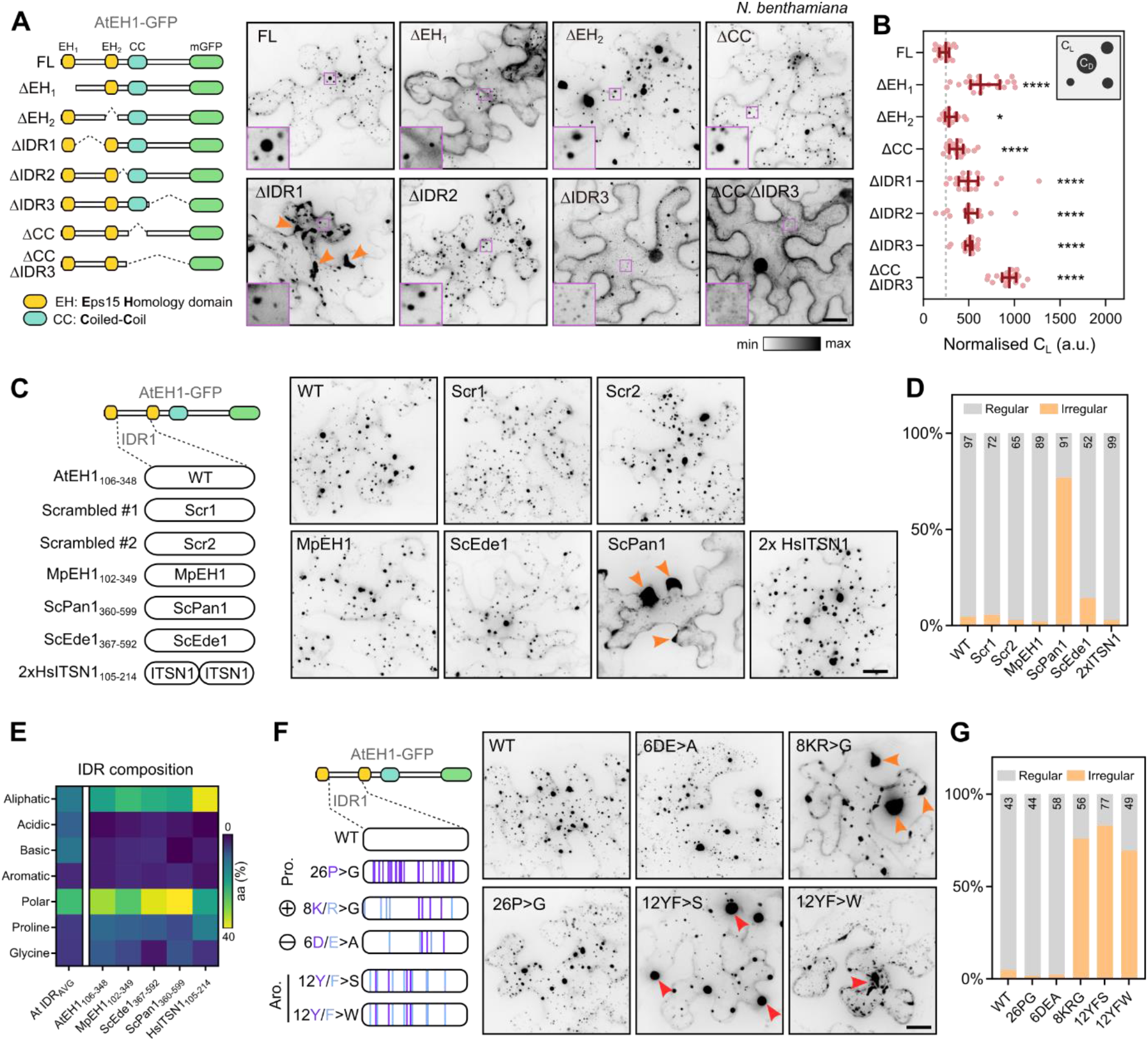
Condensation of AtEH1 is controlled by multiple regions, including IDR1. (A-B) Schematic of AtEH1 truncation constructs (UBQ10:AtEH1_domain_-GFP) and localisation in *N. benthamiana* (A). Insets show zoom in of condensates (dense phase; C_D_) and cytosol (light phase; C_L_). (B) Quantification of cytosolic protein concentration (light phase; C_L_). Bars indicate median ± 95% CI. Higher values indicate a reduced ability to phase separate. Statistics indicate significance to FL; *p < 0.05, ****p <0.0001, unpaired t-test with Welch’s correction. (C-D) IDR swap experiment. The IDR1 of AtEH1 was scrambled (Scr) or replaced with an equivalent IDR from AtEH1 homologs from moss (*Marchantia polymorpha*; MpEH1), yeast (*Saccharomyces cerevisiae*; ScPan1, ScEde1), and human (*Homo sapiens*; HsITSN1). Images of reporters in *N. benthamiana* are shown. Orange arrowheads indicate irregular condensates appearing in cell lobes. (D) Quantification of condensate distribution in cells. Number indicates sample size. (E) Heat map of amino acid composition of IDRs from the *A. thaliana* IDR proteome, and from the IDR1 of AtEH1 and equivalent sequences from related proteins. AtEH1 homologs are more similar to each other than to the average *A.thaliana* IDR sequence. Notably, ScPan1 has relatively few basic residues. (F-G) AtEH1 IDR1 mutation experiment. Proline, basic, acidic, and aromatic residues of the IDR1 of AtEH1 were mutated. Images show localisation of reporters in *N. benthamiana*. Orange arrowheads indicate irregular condensates appearing in cell lobes, red arrowheads indicate condensates with abnormal morphology. (G) Quantification of condensate distribution in cells. Number indicates sample size. Scale bars = 20 μm. **See also Figure S2.**

To investigate the function of AtEH1 IDRs, we first conducted a sequence alignment of 128 EH/Pan1 homologs throughout 700 million years of evolution. Conservation analysis revealed that IDR1 and IDR2 are relatively divergent at the single amino acid level (Figure S2B). Conversely IDR3, which contains putative short linear motifs that may facilitate protein interactions, had higher sequence conservation. Furthermore, analysis of homologous IDR1 sequences indicated that the proportion of amino acid classes was relatively conserved among plant EH/Pan1 homologs (Figure. S2C). Therefore, we infer that the sequence composition, but not the order of residues in IDR1 is functionally relevant. To test this, we scrambled the IDR1 sequence of AtEH1, or replaced it with a corresponding IDR sequence from moss (*Marchantia polymorpha*; MpEH1), yeast (*Saccharomyces cerevisiae*; ScPan1, ScEde1), or humans (*Homo sapiens*; HsITSN1). Scrambling, or replacement with MpEH1, HsITSN1, or ScEde1 IDRs resulted in cells with mostly regularly distributed condensates (Figures 2C and D). However, replacement with ScPan1 IDR caused irregularly distributed condensates which frequently accumulated in cell lobes (Figures 2C). Sequence composition analysis revealed that ScPan1 had an unusually low proportion of basic residues (1.7%), compared to AtEH1 (4.9%) (Figures 2E). These results suggest that the amino acid composition of IDR1 is critical for condensate formation.

To investigate the influence of specific residues on condensate properties, we mutated aromatic, basic, acidic, and proline residues in the IDR1 of AtEH1. Exchanging tyrosine and phenylalanine residues for tryptophan (12YF>W), or serine (12YF>S) to increase or decreasing aromatic interaction strength respectively, caused the formation of abnormally large or unusually shaped condensates (Figures 2F and G). Mutation of proline (26P>G), or acidic (6DE>A) residues had no visible effect, while mutation of basic (8KR>G) residues caused an aberrant condensate accumulation in cell lobes similar to those observed with the IDR of ScPan1. Taken together, our results show that the composition, rather than the sequence order of residues is important for controlling AtEH1 condensate properties, and that there is likely selective pressure to maintain the IDR sequence composition within an optimal range.

### AtEH1 condensates are nucleated through interactions with anionic phospholipids on the plasma membrane

Membrane surfaces have been shown to dramatically lower the concentration threshold required for protein condensates to form (Snead and Gladfelter, 2019). As AtEH1 is recruited to the plasma membrane during the early phase of endocytosis simultaneously with other TPC subunits (Wang et al., 2020), we reasoned that the plasma membrane could act as a surface to preferentially nucleate AtEH1 condensates. Supporting this hypothesis, we observed individual nucleation events on the plasma membrane in *N. benthamiana*, with condensates gradually increasing in size before dissociating from the membrane (Figure 3A and Video S1). These nucleation events had intensity profiles inconsistent with endocytic foci (Johnson et al., 2021; Narasimhan et al., 2020), and likely represent the transient accumulation of AtEH1 molecules on the membrane in this system. Condensate movement can be explained by the interaction of AtEH1 with actin (Wang et al., 2019), as treatment with the actin polymerising inhibitor Latrunculin B abolished condensate movement (Figure S3A). Furthermore, without IDR3, AtEH1 formed evenly spaced, immobile membrane-associated condensates which did not appear to coarsen (Figures 3B and 3C). This finding is consistent with reports that membrane tethered condensates are restricted in size and rapidly arrest (Snead et al., 2022). Together, these data indicate that the plasma membrane acts as a surface to nucleate AtEH1 condensates.

**Figure 3.**
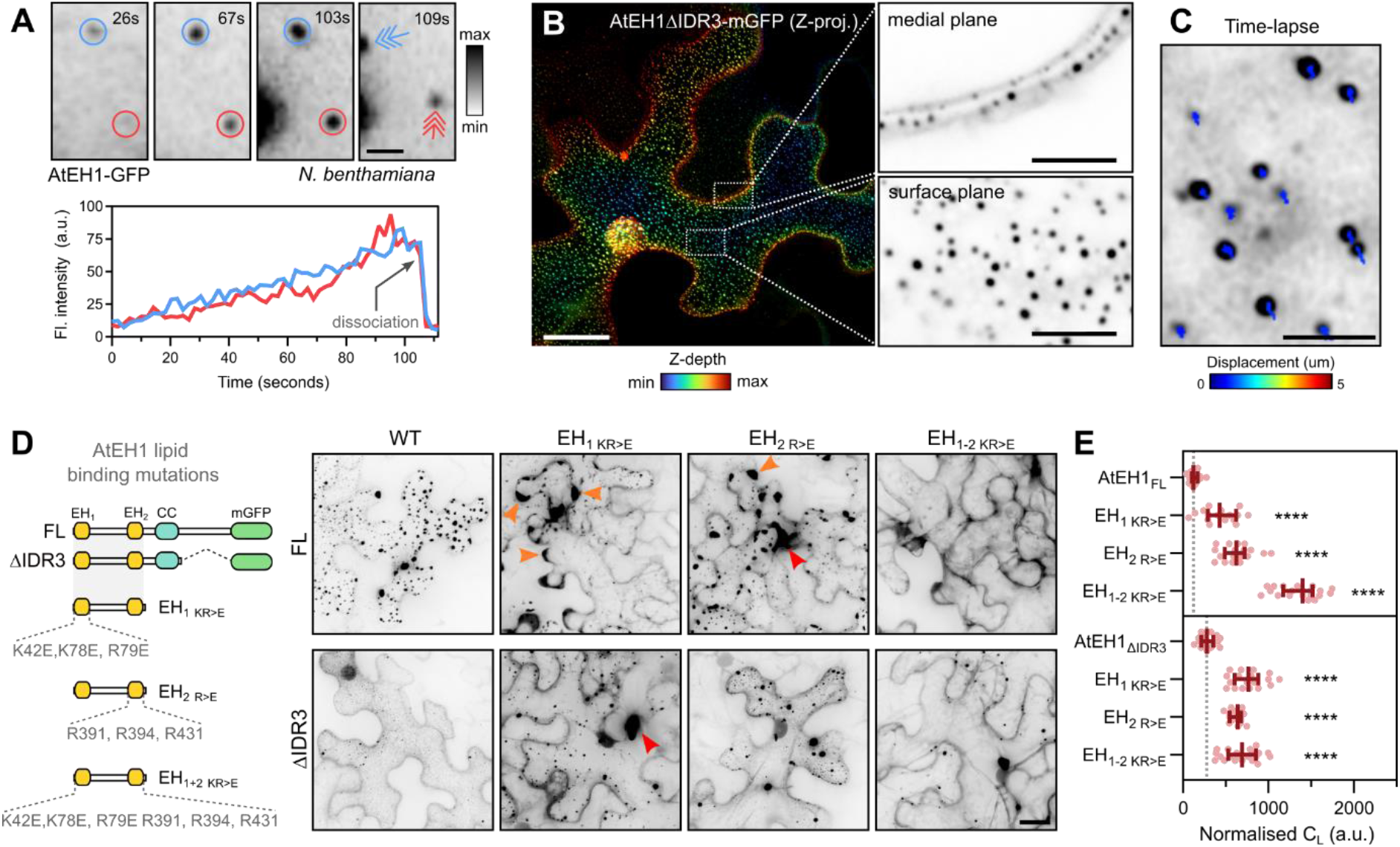
AtEH1 condensates nucleate on the plasma membrane via phospholipid binding domains. (A) Time-lapse imaging of UBQ10:AtEH1-GFP in *N. benthamiana* epidermal cells. Condensates appear as immobile plasma membrane associated puncta which gradually increase in intensity, before dissociating from their original location (arrows). (B-C) Depth colour-coded projections of UBQ10:AtEH1_ΔIDR3_-mGFP in *N. benthamiana* epidermal cells (B). Insets (single Z-plane sections) show that the condensates are restricted to the plasma membrane. Time-lapse and tracking of condensates (C). Colours indicate the total displacement of tracked condensates over a 1-minute duration. (D-E) Schematic of EH domain lipid binding mutants in AtEH1_FL_ and AtEH1_ΔIDR3_ constructs. Altered condensate distribution and properties are indicated (orange arrowheads). Condensates were not observed in AtEH1_FL_ when both EH domains were mutated. (E) Quantification of light phase (C_L_). Data is median ± 95% CI. Statistics indicates significance to the control (AtEH1_FL_ or AtEH1_ΔIDR3_), **** = p <0.0001, unpaired t-test with Welch’s correction. Scale bars = 2 μm (A), 20 μm (B, C), 5 μm (B; inset). **See also Figure S3; Video S1.**

As AtEH proteins bind negatively charged anionic phospholipids via their EH domains (Yperman et al., 2021a), we asked whether AtEH1 condensates could be nucleated by interaction with anionic phospholipids on the plasma membrane. To test this, we mutated the three positively charged arginine and lysine residues that mediate lipid binding in each individual EH domain (Yperman et al., 2021a), and examined the effect on condensate formation using full length (AtEH1_FL_) and truncated (AtEH1_ΔIDR3_) reporters (Figure 3D). Mutation of individual EH domains significantly increased the cytosolic protein concentration and lead to the formation of irregular condensates (Figures 3D and 3E), indicating that functional lipid interactions are required for normal condensate formation and properties. Moreover, combined mutation of both EH domains completely abolished condensate formation in the AtEH1_FL_ reporter, although a few condensates were observed in the truncated AtEH1_ΔIDR3_ reporter. Thus, AtEH1 condensates are primarily nucleated via phospholipid interactions, while additional cues, such as membrane-bound protein partners may act as additional nucleation factors.

Endocytosis requires the activity of multiple anionic phospholipids in the plasma membrane (Kaksonen and Roux, 2018). In plants, this includes phosphatidic acid (PA), phosphatidylinositol 4-phosphate (PI4P), and phosphatidylinositol 4,5-bisphosphate (PI(4,5)P_2_) (Noack and Jaillais, 2017; Noack and Jaillais, 2020). To determine whether specific lipids are required for condensate nucleation expressed AtEH1 in *Nicotiana tabacum* pollen tubes which maintain a distinctive lipid signature (Figure S3B) (Potocký et al., 2014). AtEH1 preferentially formed condensates at the sub-apical region of growing pollen tubes, which correlated with PA and PI4P biosensors, but not with PI(4,5)P_2_ (Figure S3B). These findings are consistent with our previous report that EH domains strongly bind PA (Yperman et al., 2021a). Furthermore, the TPC components TPLATE and TML show similar localisation patterns in pollen tubes when expressed at endogenous levels (Gadeyne et al., 2014; Van Damme et al., 2006), suggesting that the lipid-dependent nucleation of AtEH1 condensates and the recruitment of TPC to membranes during endocytosis are analogous events.

### AtEH1 is a scaffold protein which selectively sequesters endocytic proteins

Condensate assembly is often driven by highly multivalent proteins, called scaffolds, which passively recruit lower valency proteins, known as clients, to form condensates (Banani et al., 2016). Since AtEH subunits are multivalent proteins which initiate condensate formation, we asked whether they act as scaffolds to selectively recruit other endocytic proteins. To identify potential client proteins, we performed proximity labelling proteomics in *A. thaliana* cell culture using AtEH1-TurboID as a bait (Figure 4A). We also used TPLATE-TurboID (Arora et al., 2020) as a control to differentiate TPC-dependent endocytic clients from proteins identified due to AtEH1 overexpression. The endocytic proteins common to both datasets include TOM-1LIKE (TOL) proteins TOL6 and TOL9, members of an ancestral ESCRT-0 complex which function as ubiquitin receptors at the plasma membrane (Figure 4B) (Blanc et al., 2009; Herman et al., 2011; Korbei et al., 2013; Moulinier-Anzola et al., 2020). We also identified late phase endocytic proteins, including dynamins (DRP1B, DRP2A, DRP2B) which mediate vesicle scission (Fujimoto et al., 2010; Konopka and Bednarek, 2008), and AUXILIN-LIKE (AUXL) proteins, which function in vesicle uncoating (Adamowski et al., 2018).

**Figure 4.**
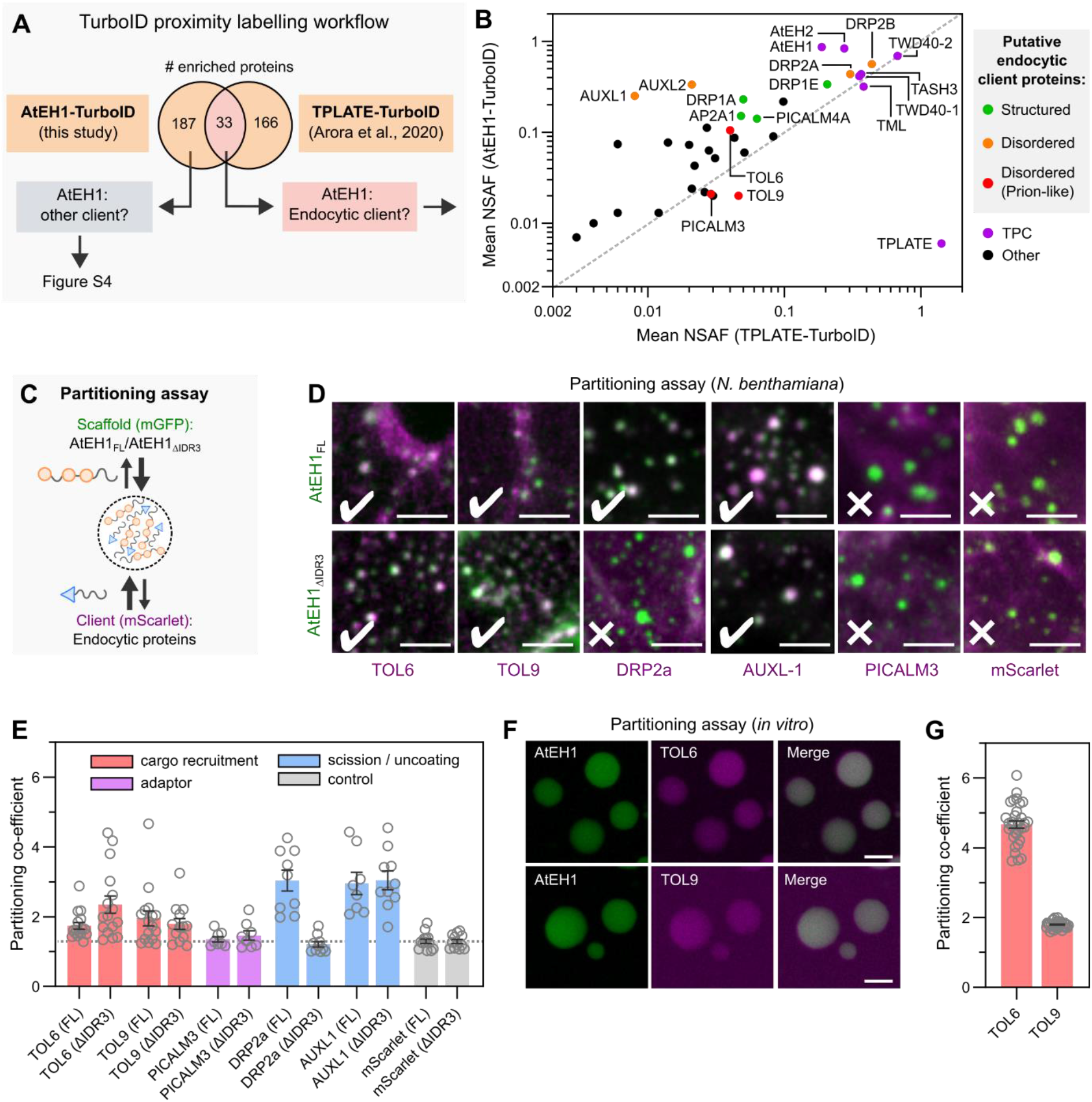
AtEH1 condensates selectively sequester endocytic machinery. (A-B) Proteomics strategy to identify endocytic client proteins of AtEH1 (A). Enriched proteins in AtEH1 and TPLATE TurboID datasets were combined, and proteins common to both datasets are plotted (B). Auxilin-like1 and Auxilin-like2 were not enriched in the TPLATE-TurboID dataset, but included due to their high enrichment and abundance in the AtEH1-TurboID dataset. Proteins which function in endocytosis are coloured based on disorder content. NSAF; normalised spectral abundance factor. (C-E) Schematic of the partitioning assay (C). Images of co-localisation of scaffolds (AtEH1_FL_-GFP and AtEH1_ΔIDR3_-GFP) with client proteins (Client-mScarlet) in *N. benthamiana* epidermal cells (D). (E) Quantification of client partitioning. Values indicates partitioning co-efficient obtained for each cell. A partitioning coefficient of 1 indicates an absence of partitioning. Client proteins are coloured according to their function in endocytosis. Bars indicate mean ± SEM. (F-G) *in vitro* partitioning assay. Purified GFP-AtEH1 and TOL6-mCherry or TOL9-mCherry were combined. TOL6-mCherry and TOL9-mCherry did not phase separate individually. (G) Quantification of client partitioning. Values indicate partitioning co-efficient from individual droplets. Bars indicate mean ± SEM. Scale bars = 5 μm. **See also Figure S4.**

We tested client recruitment using a partitioning assay in *N. benthamiana* (Figure 4C), focusing on client proteins containing disordered regions which could potentially mediate weak-multivalent interactions with AtEH1 (Figures S4A and 4B). TOL6, TOL9 and AUXL1 readily partitioned into AtEH1 condensates in *N. benthamiana*, independently of the IDR3. Conversely, DRP2a required IDR3 for partitioning (Figures 4D and 4E), indicating that IDR3 can facilitate protein interactions. PICALM3 did not partition, and may require additional factors, such as membrane curvature (Sochacki et al., 2017). Additionally, we demonstrated a direct interaction between AtEH1 and TOL6/TOL9 by partitioning assays *in vitro* (Figures 4F and 4G).

We next asked if we could identify the sequence features of TOL6 which facilitate its interaction with AtEH1. Intriguingly, we found that the AtEH1-TurboID dataset was enriched with disordered proteins, specifically proteins containing prion-like domains such as TOL6 (Figures S6A-6B). This suggests that AtEH1 mediates weak, multivalent interactions with specific prion-like domains. Supporting this, we observed partitioning of nuclear prion-like proteins identified as highly enriched in the AtEH1-TurboID dataset (Figures S4C). To specifically test whether the prion-like domain of TOL6 interacts with AtEH1, we performed a chimera experiment by transplanting the prion-like domain (PrLD) of TOL6 onto TOL3, which lacks a prion domain and does not interact with AtEH1 (Figures S4D). This TOL3-PrLD^TOL6^ chimera readily partitioned into AtEH1 condensates (Figures S4D), showing that prion-like domain recruitment is one mechanism facilitating partitioning. Collectively, our data demonstrate that AtEH1 is an endocytic scaffold which selectively recruits cytosolic adaptor proteins throughout endocytosis via multivalent interactions.

### AtEH1 condensates facilitate clathrin recruitment and re-arrangement

Endocytosis involves the ordered assembly of cytosolic adaptors and the coat protein clathrin to basket-like assemblies on the plasma membrane. Because condensation of AtEH1 is sufficient for the recruitment of multiple cytosolic endocytic accessory proteins, we asked if we could observe structural arrangements within AtEH1 condensates. To achieve this, we combined correlative light and electron microscopy (CLEM) with electron tomography (ET) to examine the ultrastructure of AtEH1-GFP condensates in a stable *A. thaliana* overexpression line (Figures S5A and S5B). We observed large cytosolic AtEH1 condensates which had remarkably similar electron density as nearby clathrin-coated vesicles (CCVs) (Figure S5C). Tomography reconstruction revealed the presence of electron dense deposits and ordered polygonal cage-like assemblies which resemble clathrin triskelia and partial clathrin cages (Figure 5A, Video S2). These observations suggest that clathrin molecules can organise into higher ordered assemblies within the liquid-like environment generated by AtEH1. We therefore examined the localisation of clathrin light chain (CLC2) in AtEH1-GFP seedlings. Similar to previous work in *N. benthamiana* (Wang et al., 2019), clathrin readily partitioned into AtEH1 condensates (Figure 5B), supporting our view that these condensates are composed of assembled clathrin molecules. Furthermore, we observed substantial clathrin accumulation around some condensates (Figure 5B), suggesting that clathrin has an inherent affinity for these assemblies. These findings show that condensation of AtEH1 is sufficient to recruit and concentrate clathrin, and facilitate its re-arrangement into ordered assemblies.

**Figure 5.**
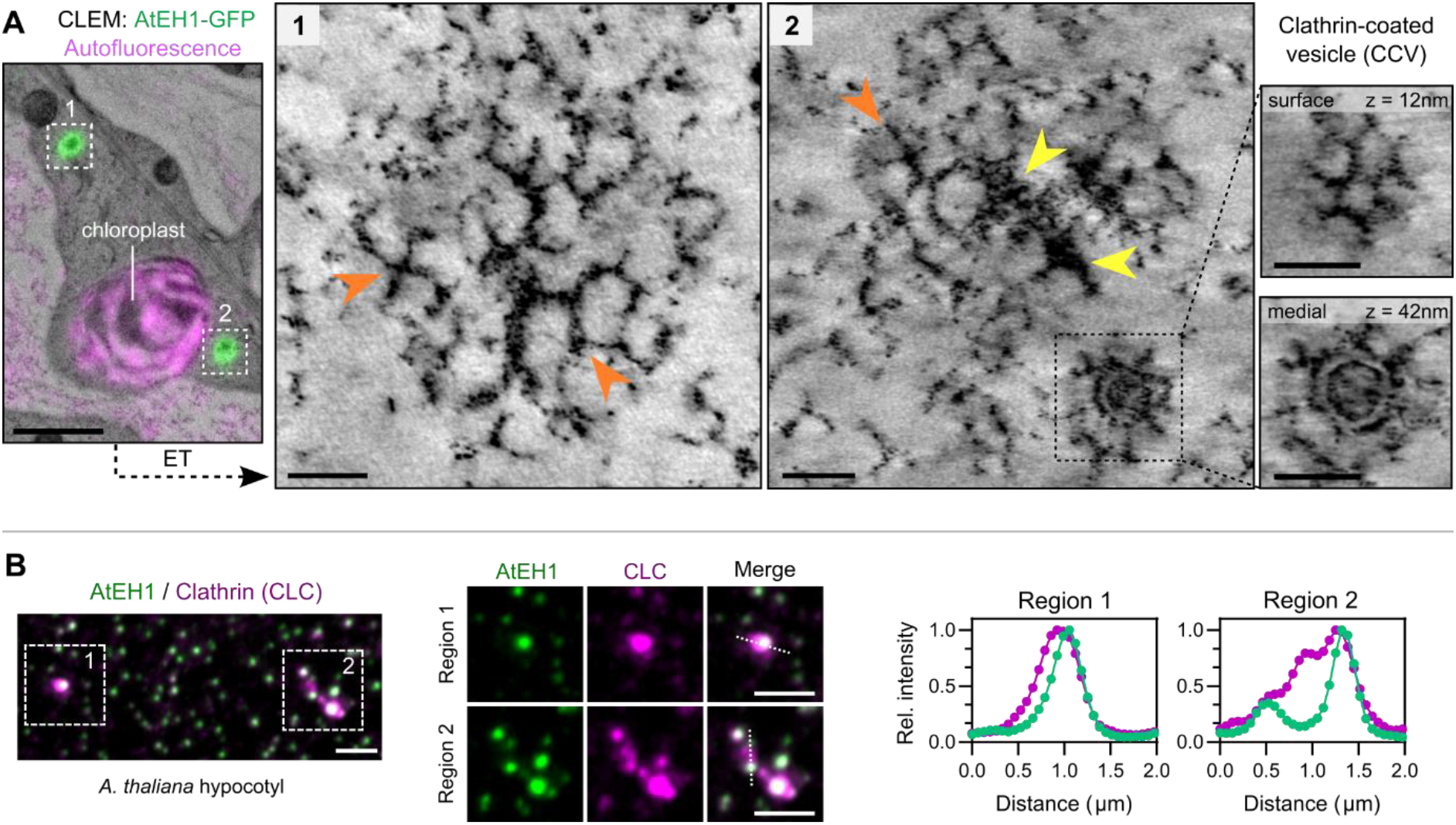
AtEH1 condensates facilitate clathrin recruitment and re-arrangement. Correlative light and electron microscopy (CLEM) combined with electron tomography (ET) of hypocotyl sections from *A. thaliana* seedlings over-expressing AtEH1 (35S:AtEH1-GFP). Clathrin-like lattices and triskelia (orange arrowheads), and electron dense accumulations (yellow arrowheads) are formed within AtEH1 condensates. Inset shows a clathrin-coated vesicle (CCV) nearby the condensate. (A) Airyscan images of clathrin light chain 2 (CLC-mKo) and AtEH1 (35S:AtEH1-GFP) in *A. thaliana* hypocotyl cells. Plot profiles show normalised fluorescence intensities from the indicated lines. Clathrin is observed within and surrounding the condensates. Scale bars = 50nm (A; ET), 2 μm (A; CLEM, B). **See also Figure S5; Video S2.**

### Concentration of TPC on a membrane is sufficient for condensate formation and clathrin assembly

Our previous work suggests TPC exists as a stable octameric complex in the cytosol, which is recruited collectively to the plasma membrane during the initiation of endocytosis (Wang et al., 2020; Yperman et al., 2021b). Further supporting this, all TPC subunits (except LOLITA) were enriched in a large scale biotinylated isoxazole (b-isox) precipitation experiment which selectively precipitates disordered proteins (Zhang et al., 2022) (Figure S6A). We independently confirmed these findings by b-isox precipitation and immunoblotting against AtEH1 (highly disordered TPC subunit) and TPLATE (low disorder TPC subunit), which showed enrichment of both proteins (Figure 6A). Furthermore, highly disordered endocytic client proteins, including TOL6, TOL9, AUXL1, AUXL2, and DRP2a were also identified as enriched in the b-isox fraction (Figure S6A), while structured proteins like clathrin, tubulin and AP-2 were not (Figure 6A and S6A).

**Figure 6.**
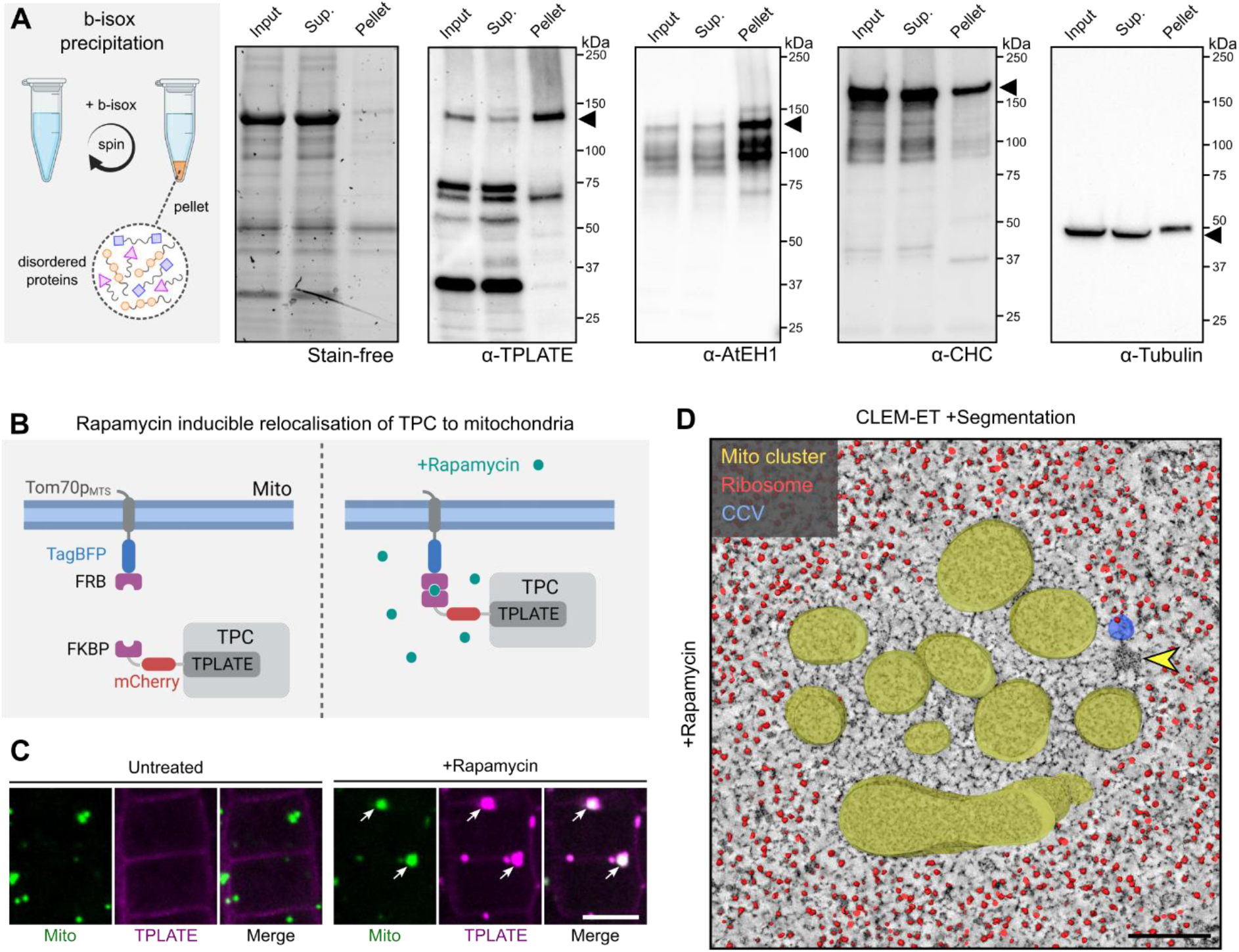
Chemical induced re-localisation of TPC to mitochondria is sufficient for condensate formation and clathrin assembly. (A) Biotinylated isoxazole (B-isox) enrichment assay. *A. thaliana* cell lysates were treated with b-isox to selectively precipitate IDR containing proteins after centrifugation (pellet fraction). The input, supernatant, and pellet fractions were used for western blots with the indicated antibodies. Stain free gel shows selective enrichment of proteins in the pellet fraction. Tubulin is used as a negative control. The experiment was repeated twice with similar results obtained. (B) Schematic of the chemically induced TPLATE mitochondria re-localisation experiment. (C) Images of *A. thaliana* root cells expressing Tom70p_MTS_-TagBFP2-FRB (Mito) and TPLATE-mCherry-FKBP (TPLATE) from untreated, and rapamycin-treated seedlings. TPLATE is re-localised to mitochondria clusters after rapamycin treatment (arrows). (D) Segmented ET reconstruction of a mitochondria cluster from a rapamycin treated *A. thaliana* root cell after localisation by CLEM (Fig. S6C). The interior of the mitochondria cluster is devoid of ribosomes, indicating a liquid-like environment. The yellow arrowhead indicates an electron dense cluster. The partial volume rendering of the reconstruction is shown; full volume segmentation in Fig. S6C. Scale bars = 10 μm (C), 200 nm (D). **See also Figure S6; Video S3 and S4.**

Our previous experiments relied on overexpression of AtEH1 to generate condensates. However, to be functionally relevant for endocytosis, condensation must occur via TPC as a whole on the membrane. We therefore asked whether the concentrated assembly of TPC at endogenous stoichiometric ratio on a membrane is sufficient to generate condensates. Because it is not possible to examine the ultrastructure of endocytic pits *in planta* due to poor membrane preservation, we used an inducible FRB-FKBP rapamycin-based interaction system to concentrate TPC on mitochondrial membranes (Figure 6B and S6B-C)(Robinson et al., 2010; Winkler et al., 2021). Using this system, we inducibly re-localised TPLATE to mitochondria clusters (Figure 6C and Video S3). These clusters contained AtEH1 and clathrin, confirming that other TPC subunits and interacting proteins were recruited (Figure S6D). Next, we employed a CLEM-ET and segmentation approach to characterise the 3D ultrastructure of these TPC-mitochondria clusters (Figure S6E). We observed distinct ribosome exclusion zones in the interior of the mitochondria clusters after rapamycin treatment, which were not observed in the untreated control (Figure 6D and S6E). This zone contained ordered, clathrin-lattice-like assemblies which spanned the entirety of the area (Figure 6D and Video S4). We also observed clathrin coated vesicles (CCVs) and electron dense accumulations on the edges of the mitochondria clusters (Figure 6D and S6E). These observations were reminiscent of our CLEM-ET ultrastructural data of condensates driven by AtEH1 alone. Together, these data indicate that concentration of TPC on a membrane is sufficient to generate condensates which promote clathrin recruitment and re-arrangement. Thus, AtEH1 and AtEH2 promote condensation as part of TPC.

### The physical properties of TPC driven condensates are important for endocytosis progression

Since TPC recruits clathrin and other endocytic machinery, we asked whether condensation of TPC is essential for functional endocytosis, for example by enabling the recruitment and re-arrangement of endocytic machinery throughout the process. We reasoned that we could manipulate the properties of TPC condensates by altering the aromatic interaction strength of the IDR1 of AtEH1, since aromatic residues have been shown to promote phase separation of prion-like domain proteins (Brangwynne et al., 2015; Martin et al., 2020). We created modified AtEH1 reporters by substituting phenylalanine and tyrosine residues with serine (12YF>S) or tryptophan (12YF>W) to decrease or increase IDR interaction strength, respectively (Figure 2F). Because AtEH1 condensates are motile, we could infer the material properties of AtEH1 IDR1_mut_ condensates by high speed 4D imaging. Surface renderings reveal that AtEH1_YF>W_ condensates were highly dynamic and irregular shaped, suggesting a gel-like material state (Figure 7A-7B, and Video S5). Comparatively, AtEH1_YF>S_ and AtEH1_WT_ condensates were more spherical, indicating a more liquid-like material state (Figure 7A-B). To further assess the molecular properties of AtEH1 IDR1 mutants, we performed a ratiometric FRAP assay by expressing both AtEH1_WT_ and AtEH1_mut_ constructs from an individual T-DNA locus, and comparing the FRAP half recovery time (t_1/2_). Recovery of AtEH1_YF>S_ was significantly faster than the AtEH1_WT_ control, indicating a decrease in intramolecular interaction strength between AtEH1 molecules (Figure 7C and 7D). Conversely, AtEH1_YF>W_ showed a delayed recovery, indicating an increase in intramolecular interaction strength. Our results demonstrate that altering the IDR1 interaction strength of AtEH1 can modulate the dynamics and material properties of AtEH1 condensates.

**Figure 7.**
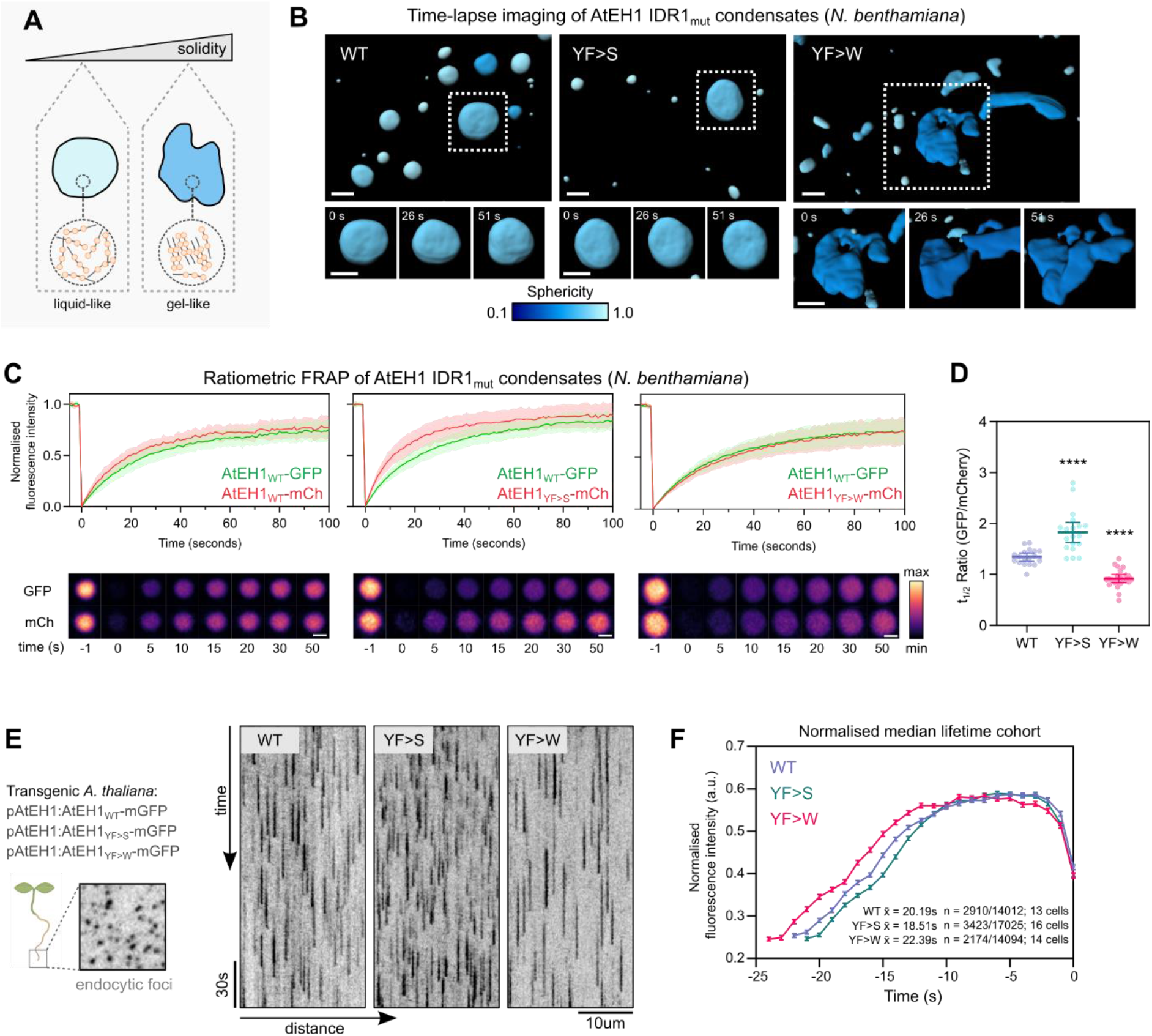
The physical properties of TPC driven condensates are important for endocytosis progression. (A-B) Schematic representation of condensate material properties (A). In gel-like condensates, proteins form more rigid interactions and have reduced molecular exchange. 3D surface rendering of condensates from *N. benthamiana* epidermal cells expressing UBQ10:AtEH1_WT/mut_-GFP constructs (B). Condensates are coloured by sphericity, with a value of 1 indicating a perfect sphere. (C-D) Ratiometric FRAP of condensates from *N. benthamiana* epidermal cells expressing two reporters from a single plasmid (C). Plot shows FRAP recovery curves from a binned subset of the data; bars indicate mean ± SD, n= 8. (D) Quantification of t_1/2_ ratio from all FRAP curves. Bars indicate mean ± 95% CI. Statistics indicate difference from WT; **** p<0.0001, unpaired t-test with Welch’s correction. (E-F) Kymograph analysis of endocytic events from root cells of *A. thaliana eh1-1* homozygous mutant lines rescued by AtEH1:AtEH1_WT/MUT_-GFP. (F) Fluorescence intensity profile calculated from the median lifetime of endocytic events; error bars indicate SEM. The number of endocytic events in the median lifetime cohort, and the total number of measured events is indicated. Scale bars = 5 μm (A), 2 μm (C), 10 μm (E). **See also Figure S7; Video S5.**

Finally, we aimed to understand how altering the properties of TPC condensates affects endocytosis. To achieve this, we generated native promoter (pAtEH1:AtEH1-mGFP) wild type (FL), and aromatic mutant (YF>S, YF>W) *A. thaliana* reporters. These lines were functional as they rescued the *A. thaliana* male sterile *eh1-1* mutant (Figure S7A and S7C). We identified multiple complementation lines for each construct containing close to endogenous levels of AtEH1 fusion protein (Figure S7B). To determine if these plants had defects in bulk endocytic flux, we assayed AtEH1_WT_ and AtEH1_mut_ *A. thaliana* seedlings using an FM4-64 dye internalisation assay (Figure S7D). Bulk internalisation was not significantly different in control AtEH1_WT_ lines to WT Col-0 seedlings, but was slightly reduced in AtEH1_YFS_ lines, and vastly reduced in AtEH1_YFW_ lines Figure S7D and S7E. Thus, alteration of condensate properties impairs endocytosis.

To test whether these defects are due to an alteration in endocytosis dynamics, we performed live-cell imaging of endocytic foci in *A. thaliana* root cells (Figure 7E). Initial kymograph analysis showed that the decreased intramolecular interaction strength in AtEH1_YFS_ seedlings corresponded to a reduction in endocytosis lifetime, while increasing intramolecular interaction strength in AtEH1_YFW_ extended endocytosis lifetime (Figure 7E). We further assessed this quantitatively using an automated analysis of endocytosis lifetime using an automated detected and analysis pipeline (Aguet et al., 2013; Johnson et al., 2020). Consistent with the kymograph analysis, analysis of >14,000 endocytic events from each genotype revealed a reduction in endocytic lifetime in AtEH1_YFS_ seedlings compared to the AtEH1_WT_ control (Figure 7F), and an increase in lifetime of AtEH1_YFW_ mutant seedlings. Thus, the molecular re-arrangement and material properties of condensates driven by AtEH1/TPC influences the underlying progression of endocytosis. The altered endocytic progression also had a physiological effect, as we observed delayed root gravitropism responses in AtEH1_YFW_ lines, a process which requires TPC-dependent endocytic activity (Gadeyne et al., 2014). Taken together, our results show that condensate physical properties modulate endocytosis and subsequently impact plant adaptive growth, and that the IDR sequences which determine these properties likely converge on an evolutionary optimal code.

## Discussion

The formation of clathrin coated pits during endocytosis requires the dynamic re-arrangement of a meshwork of proteins including clathrin, adaptors, and accessory proteins (Chen and Schmid, 2020; Schmid and McMahon, 2007). Here, we discovered that AtEH proteins drive biomolecular condensation of the TPLATE complex during clathrin-mediated endocytosis in plants. We show that these condensates enable the recruitment and assembly of clathrin and accessory proteins, from the initiation of endocytosis to vesicle scission. The biophysical properties of these condensates are partially derived from the sequence chemistry of the IDR of the AtEH proteins, which share strong similarity with their human and yeast counterparts. We demonstrate *in vivo,* that alteration of condensate properties impairs the basic progression of endocytosis and adaptive plant growth. Therefore, biomolecular condensation is an essential biophysical mechanism which can help explain how collective interactions between endocytic machinery generate a clathrin coated pit.

### Plasma membrane recruitment nucleates AtEH1/TPC condensation

Our work highlights a link between the nucleation of TPC condensates on the membrane and the initiation of endocytosis. Our *in vivo* CLEM data shows that concentration of TPC on a membrane is sufficient to trigger condensation, likely by limiting molecular diffusion to a two-dimensional plane which lowers the nucleation energy barrier (Snead and Gladfelter, 2019; Snead et al., 2022). In our model, TPC is initially recruited to the membrane by direct interactions with phospholipids via AtEH subunits. These initial assemblies are self-reinforced by recruitment of additional TPC units through multivalent interactions mediated by AtEH subunits. Once TPC is sufficiently concentrated on the membrane, this promotes the downstream recruitment of adaptors and clathrin to initiate endocytosis and the assembly of clathrin-coated pits. In this way condensation functions as an organisational hub (Shin and Brangwynne, 2017). This model is consistent with work in humans and yeast, where initiator proteins Eps15/FCHo and ScEde1 promote condensation and recruitment of endocytic accessory proteins to the membrane via interactions with the phospholipid PI(4,5)P_2_ (Day et al., 2021; Kozak and Kaksonen, 2022).

How does condensation of TPC promote endocytosis initiation at specific sites? AtEH1 remains at the plasma membrane upon destabilisation of TPC (Wang et al., 2021), indicating a high affinity for membranes. Specifically, AtEH1 interacts with PA and phosphoinositides (Yperman et al., 2021a), two key phospholipids implicated in endocytosis in plants (Noack and Jaillais, 2020; Platre et al., 2018). These lipids may act as a molecular signature to specify the recruitment and condensation of TPC. Interaction of TPC with PI4P via the muniscin-like subunit TML may further enhance recruitment specificity (Yperman et al., 2021b), or could promote complex stability at the membrane, similar to the role of FCHo (Bhave et al., 2020; Henne et al., 2010; Lehmann et al., 2019). We also identified a link between AtEH1 and TOM1-LIKE (TOL) proteins, which function in plants as ubiquitin receptors, analogous to the ESCRT-0 complex (Korbei et al., 2013; Mosesso et al., 2019; Moulinier-Anzola et al., 2020). We speculate that recognition of TOL proteins by AtEH subunits promote condensation and endocytosis initiation at membrane regions enriched for ubiquitinated cargo, possibly in co-operation with the TPC subunit TASH3 (Grones et al., 2022). This coupling of TPC recruitment and condensation to the co-operative recognition of lipids and ubiquitinated cargo would ensure the highly regulated assembly of clathrin-coated vesicles on the plasma membrane. Similar, lipid— or receptor—mediated nucleation mechanisms may exist to drive the selective assembly of other membrane associated condensates, for example during autophagy (Fujioka et al., 2020). However, direct observations of nucleation events on the membrane *in vivo* remains challenging, owing to the transient and small size of such assemblies, especially during endocytosis (Cocucci et al., 2012).

### Condensation of TPC promotes clathrin and accessory protein assembly

During the dynamic growth phase of clathrin coated pits, clathrin molecules are rapidly deposited at the edge of the growing lattice (Robinson, 2015; Sochacki and Taraska, 2019). Eps15, FCHO, Intersectin and ScEde1 partition to the lattice edge, and are continuously displaced from entering the inner lattice (Mund et al., 2018; Partlow et al., 2022; Sochacki et al., 2017). A similar model has been proposed with TPC confined to the lattice edge in plants (Johnson et al., 2021). Our CLEM-ET experiments reveal that condensation of AtEH1/TPC promotes the recruitment and re-arrangement of clathrin molecules in a liquid-like environment.

TPC is well positioned to catalyse the sustained assembly of clathrin molecules into the growing lattice, and could explain why TPC arrives before clathrin on the membrane (Gadeyne et al., 2014; Narasimhan et al., 2020). How clathrin is recruited into these condensates is not clear, but this could involve direct interaction with TPC subunits (Gadeyne et al., 2014; Van Damme et al., 2011). However, these interactions alone are not sufficient for the co-purification of clathrin with TPC in b-isox precipitation experiments. Clathrin recruitment and assembly in condensates is therefore likely driven by a combination of transient, low-affinity interactions, and relatively stable, high-affinity interactions between clathrin, TPC, and other clathrin-interacting molecules, such as AP-2. These collective interactions may occur within an ideal physiochemical generated by condensation of TPC, resulting in the efficient assembly of clathrin at the edge of growing lattices. Furthermore, condensation could help explain mechanistically how clathrin is rapidly turned over throughout endocytosis (Avinoam et al., 2015), which is a key requirement to build an expanding clathrin cage.

Besides clathrin, our data suggest that TPC facilitates the assembly and re-arrangement of dynamin and auxilin, likely at the neck of invaginated clathrin coated pits. Condensation of dynamin has been shown to enhance its scission activity (Imoto et al., 2022), supporting our model that condensation enables TPC to act as an organisational hub to assemble cytosolic endocytic proteins throughout endocytosis. Our work also highlights evolutionary differences between endocytosis across kingdoms, as condensation of ScEde1 and Eps15-FCHo is coupled to endocytosis initiation, as these components are disassembled from the membrane before scission (Henne et al., 2010). Given that middle-late arriving proteins ScSla1 (Bergeron-Sandoval et al., 2021), and human endophilin can phase separate (Mondal et al., 2022), condensation throughout endocytosis in these systems may be driven by discrete modules, rather than a stable complex.

### IDR composition is a hidden regulator of endocytic dynamics

The physical properties of condensates are controlled by multivalent, co-operative interactions which form a physically cross-linked network (Boeynaems et al., 2018). Our data indicates that the sequence composition of the IDR1 of AtEH1 plays a significant role in controlling the properties of TPC driven condensates. Modification of IDR1 interaction strength influences the material state and molecular exchange rate of these condensates, presumably by affecting the degree of cross-linking between AtEH1 molecules. Importantly, we show that divergence from this evolutionarily ideal composition reduces endocytosis efficiency *in vivo*. Given that the IDR composition of AtEH/ homologs is similar throughout evolution, there is likely an optimal physiochemical environment derived from IDR properties which provide a balance between ordered, solid-like, and flexible, liquid-like network assemblies. Notably, artificially strengthening Eps15 dimerisation during early clathrin coated pit formation impairs endocytosis in human cells (Day et al., 2021), highlighting that multivalency driven by scaffold dimerisation or IDR interactions is an essential aspect of endocytosis.

It is unclear how the biophysical properties and selectivity of protein interactions are encoded into the sequence chemistry of the IDRs of AtEH proteins (Choi et al., 2020; Martin and Mittag, 2018). Furthermore, whether these IDR-derived condensate properties are unique to endocytosis, and how they have been evolutionarily tuned is unknown. Given that AtEH1 and AtEH2 are not redundant, and most land plants contain at least two EH/Pan1 homologs, independent proteins may mediate differential protein recruitment to reprogram endocytosis in an environmentally responsive manner.

In summary, our study identifies biomolecular condensation as a fundamental organisation principle promoting the assembly of clathrin and other key molecules throughout endocytosis. TPC functions as a central organiser for endocytosis in plants, and achieves this through using core principles which appear common between eukaryotic organisms. Our findings have implications for understanding how condensates form on membranes, and how they act as an organisation hub to generate selective and dynamic interaction networks to facilitate vesicle trafficking.

## Supporting information

membrane associated condensate formation video

AtEH1 condensate ET video

knocksideways in Arabidopsis root video

segmented ET video Arabidopsis root

condensate dynamics imaris rendering

list of constructs used in the manuscript

## Acknowledgements

This work was supported by the European Research Council Grant T-REX 682436 (D.V.D.), 852136 (A.B.); the Research Foundation– Flanders (FWO) 1226420N (P.G.), 12S7222N (J.M.D.), 1124621N (A.D.M.), an EMBO Scientific exchange grant 9253 (J.M.D); Czech Science Foundation grant 22-35680M (R.P.), National Natural Science Foundation of China grant 32161133001 (X.F.) and Beijing Natural Science Foundation grant JQ21020 (X.F.). We acknowledge the Imaging Facility of the Institute of Experimental Botany AS CR supported by the MEYS CR (LM2018129 Czech-BioImaging) and IEB AS CR. We acknowledge the VIB BioImaging Core, and Anneke Kremer for help with the Amira analysis. We thank Evelien Mylle for technical support.

## Author contributions

J.M.D initiated the project, designed, and performed all experiments unless otherwise indicated. Y.W. purified proteins, performed in-vitro assays and yeast localisation. L.B., C.C. and J.M.D performed the CLEM-ET experiments. A.D.M. performed tobacco partitioning assays. R.H., R.P. and M.P. performed the pollen tube experiments. R.P. made the EH sequence alignment and phylogenetic tree. M.V. and P.P. cloned constructs. P.G. performed the immunolocalisation assay. D.E. analysed proteomics data. R.H. and M.B. performed image analysis. J.W. generated the rapamycin-based plant lines. N.S. performed rapamycin time-lapse imaging. M.F., A.B., G.D.J., R.P., X.F., and D.V.D supervised research. J.M.D and D.V.D wrote the manuscript. All authors contributed to finalising the text.

## Materials and Methods

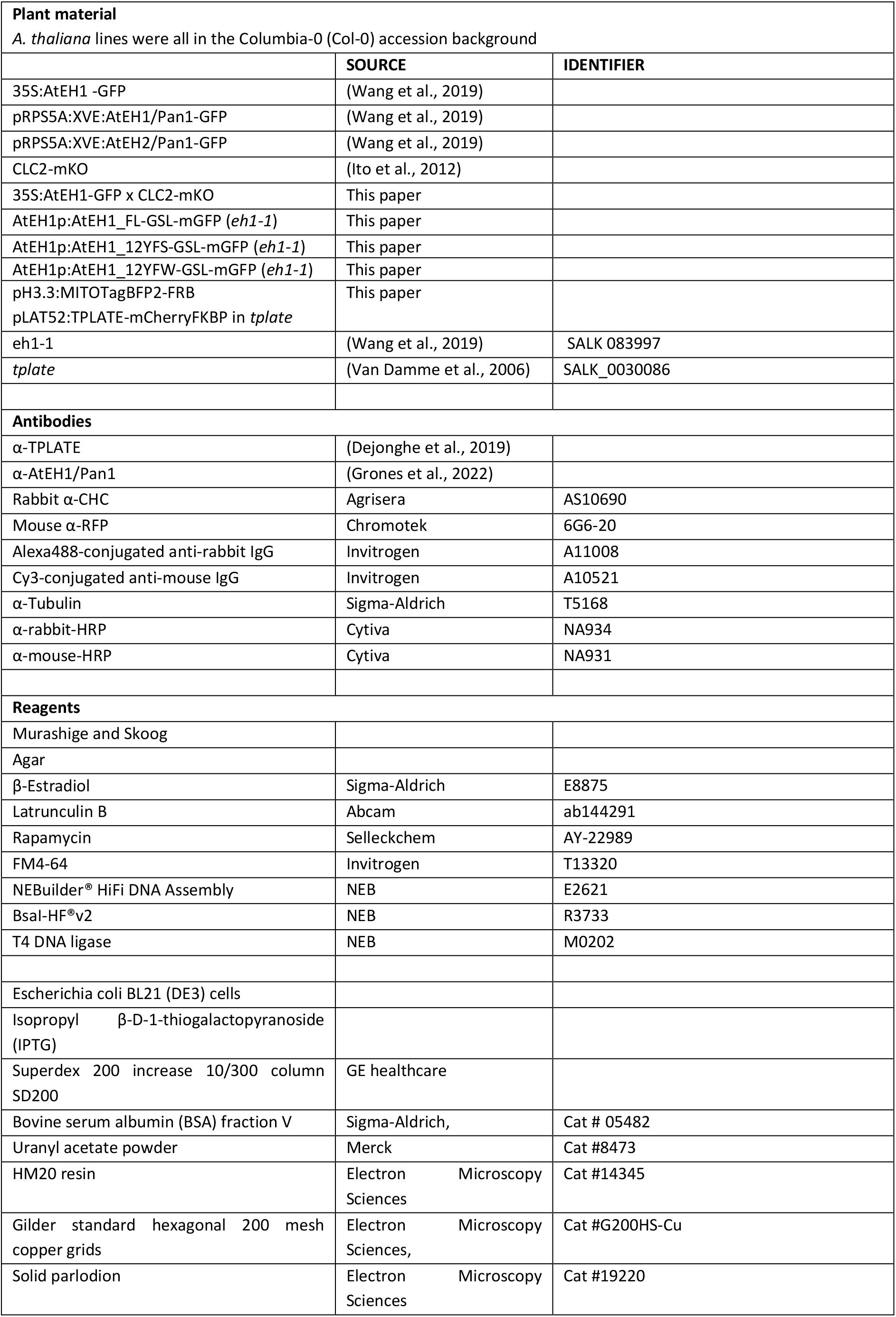

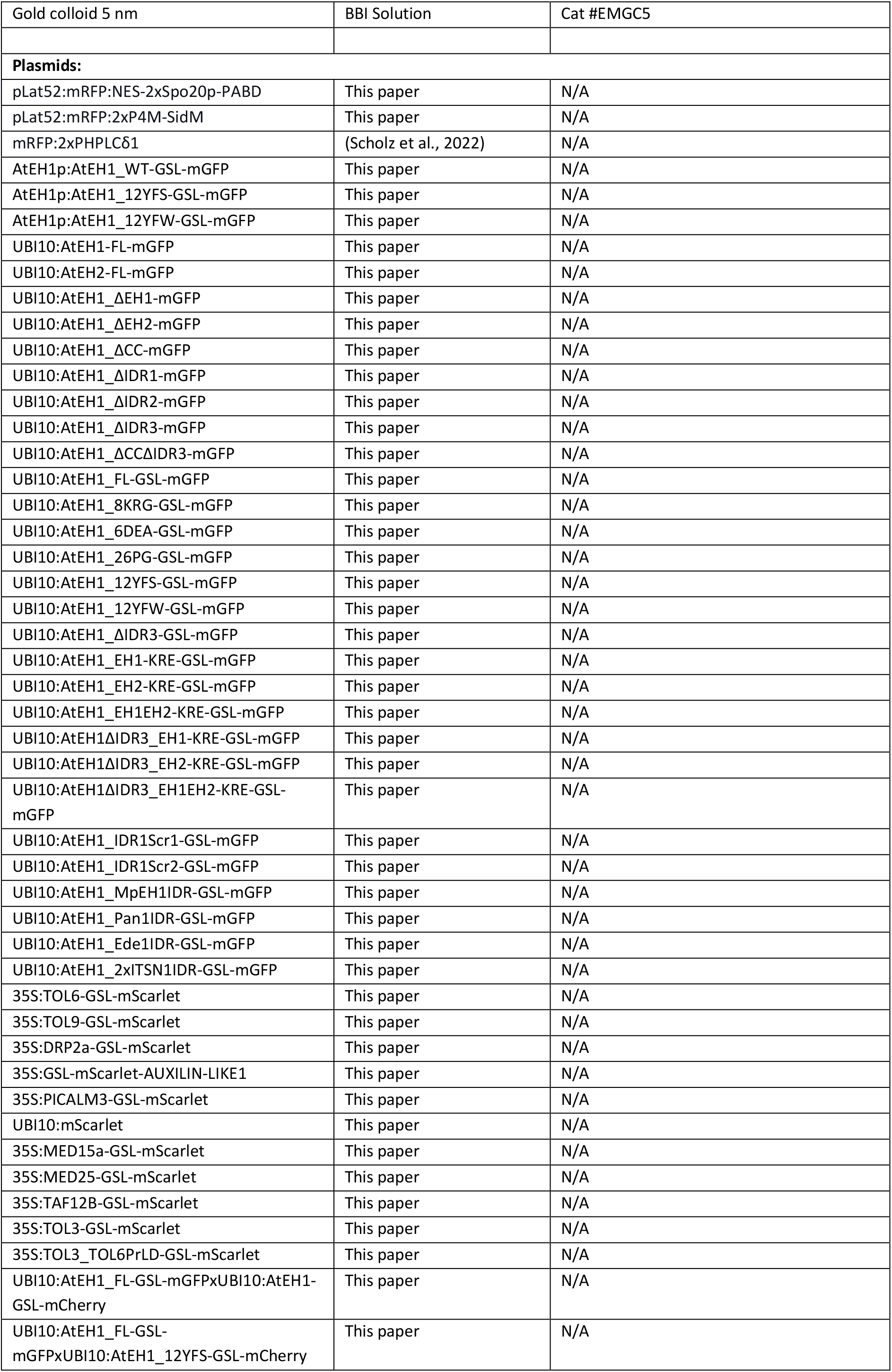

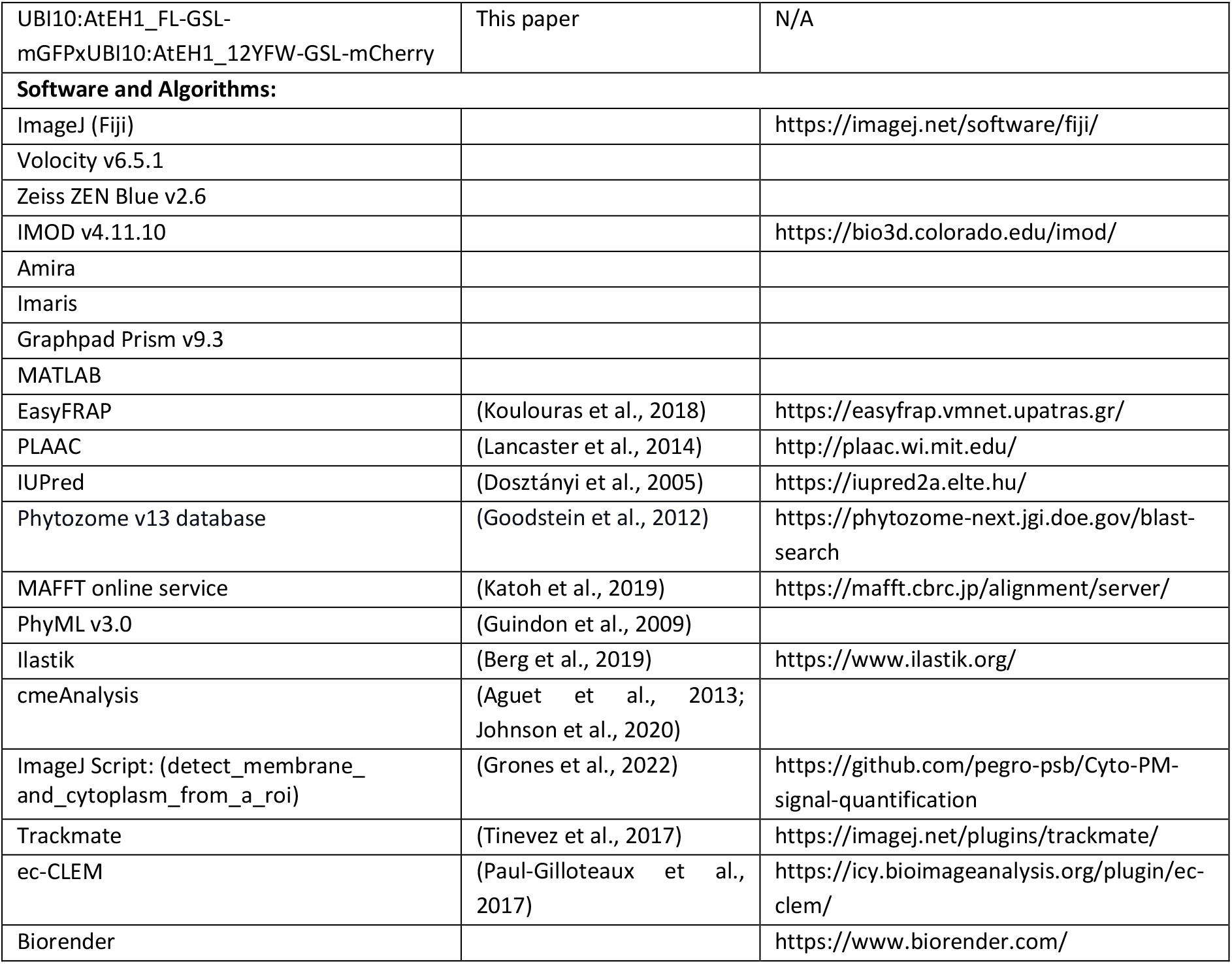

### Plant material and growth conditions

*Arabidopsis thaliana* accession Columbia-0 (Col-0) plants were used for all experiments. Seeds were surface sterilised by chlorine gas and grown on ½ strength Murashige and Skoog (½ MS) medium containing 0.6% (w/v) agar, pH 5.8, without sucrose. Seedlings were stratified for 48 h at 4°C in the dark, and transferred to continuous light conditions (68 μE m−2 s−1 photosynthetically active radiation) at 21 °C in a growth chamber. Imaging was performed on 4–5-day old seedlings unless otherwise indicated. β-Estradiol induction of the pRPS5A:XVE lines was performed by transferring 3– day-old seedlings to ½ MS medium containing 1 µM β-estradiol.

### Chemical treatments

β-Estradiol (20 mM stock in DMSO), Latrunculin B (4 mM stock in DMSO), and Rapamycin (10mM stock in DMSO) were used at the concentrations indicated.

### Molecular cloning

All constructs were cloned and assembled using the Green Gate cloning system unless otherwise indicated (Lampropoulos et al., 2013), and are detailed in Table S1. Entry vectors were assembled by Gibson assembly using NEBuilder® HiFi DNA Assembly. For generation of mutated constructs, gene fragments were synthesised using Twist Bioscience.

To generate the constructs used for heterologous expression in yeast cells, AtEH1 and its variants coding sequences were amplified and cloned into pDUAL-Pnmt1-yeGFP vector by the ClonExpress II One Step Cloning Kit (Vazyme, C112). The constructs for *in vitro* protein expression were cloned by inserting AtEH1(FL), AtEH1(ΔIDR3) and AtEH1(ΔCCΔIDR3) coding sequences into MBP-HIS-GFP vector, with maltose binding protein (MBP) at N-terminus following a TEV cleavage site and a green fluorescence protein (GFP) at C-terminus. The sequences of TOL6 and TOL9 were amplified and inserted into the pRSFduet-6×His-mCherry vector to generate mCherry-TOL6 and mCherry-TOL9 constructs for *in vitro* protein expression.

To prepare the PA marker construct (pLat52::mRFP:NES-2xSpo20p-PABD), NES-Spo20p-PABD was first amplified using P1 and P2 primers and YFP:Spo20p-PABD (Potocký et al., 2014) as a template and introduced into mRFP:2xSpo20p-PABD (Pejchar et al., 2020) using XbaI/SpeI sites to make mRFP:NES-2xSpo20p-PABD. To prepare the PI4P marker construct (pLat52::mRFP:2xP4M-SidM), 2xP4M-SidM was first amplified using P3 and P4 primers and GFP:P4M-SidM2x (a gift from Prof. Tamas Balla, Addgene plasmid # 51472) as a template and introduced into the vector pHD22, kindly provided by Prof. Benedikt Kost, using XbaI/ApaI sites to make mRFP:2xP4M-SidM. The construct for the PI(4,5)P_2_ marker, mRFP:2xPHPLCẟ1, was described previously (Scholz et al., 2022).

### Generation of A. thaliana reporter lines

To generate the pEH1:EH1-GFP reporters, 2356 bp promoter and 5’UTR were fused to the CDS of AtEH1 and mGFP using Golden Gate cloning. Transformed via floral dip (Clough and Bent, 1998) into plants heterozygous for the *eh1-1* T-DNA (SALK 083997). Positive transformants were selected by spraying with Basta solution on seedlings grown in soil. Heterozygous *eh1-1* plants were selected, and in the following generation seedlings homozygous for *eh1-1* were identified by genotyping and confirmed via Western blot. To generate the pH3.3-MITOTagBFP2-FRB pLAT52-TPLATE-mCherryFKBP *tplate* line, plants heterozygous for the *tplate* T-DNA (SALK 0030086) were transformed with pLAT52-TPLATE-mCherryFKBP by floral dip. T1 Plants were selected with 25 mg/L hygromycin for presence of pLAT52::TPLATE-mCherry-FKBP, and genotyped to select *tplate(+/-)*. T2 plants were screened to identify *tplate* (-/-) plants. The following T3 seedlings were transformed with pH3.3::MITO-TagBFP2-FRB* (Winkler et al., 2021). T1 plants were selected with 10 mg/L Basta, and plants homozygous for pH3.3-MITOTagBFP2-FRB, pLAT52-TPLATE-mCherryFKBP, and *tplate* were identified in the following generations.

### Gravitropism assay

5 day old *A. thaliana* seedlings were grown on ½ MS media. Seedlings of similar sizes were first transferred to a fresh ½ MS plate to reduce variability due to potential germination rate differences, and then gravistimulated by turning the plate 90°. Images were acquired every 15 minutes using a Canon EOS camera. The time point of 8 hours was chosen for analysis as this is when we observed near complete bending of WT Col-0 plants. The root tip angle was calculated in Fiji using the angle tool, and the results were placed into 22.5° bins.

### Transient expression in Tobacco epidermis

*N. benthamiana* plants were grown in a greenhouse under long-day conditions (6–22 h light, 100 PAR, 21 °C) in soil (Saniflor osmocote pro NPK: 16-11-10 + magnesium and trace elements). Transient expression was performed by leaf infiltration according to (Sparkes et al., 2006) using GV3101 Agrobacterium strains containing plasmids of interest. To reduce silencing, an agrobacterium strain containing the p19 silencing inhibitor was co-infiltrated for all experiments. Transiently transformed *N. benthamiana* were imaged three to four days after infiltration.

### Pollen tube transformation, imaging, and analysis

Experiments were performed step by step according to the protocol described in (Noack et al., 2019). Transformed tobacco (*Nicotiana tabacum*) pollen grains were observed 6 hours after transformation with a Zeiss LSM900 laser scanning microscope (Axio Observer 7, inverted) with Airyscan 2 multiplex 4Y mode. A 40x W 1.2 NA Plan Apochromat objective was used during image acquisition. Images were acquired in ZEN blue using the smart set up configured the light paths for TagYFP and mCherry. Time-lapse of tobacco pollen tubes was acquired for 30-90s.

Analysis of plasma membrane intensities was performed in Fiji. A region of interest (ROI) was selected with the “Segmented line” tool over the plasma membrane region of the middle section of the pollen tube for each time point. For each time point, the AtEH1 (green) and biosensor (red) channel intensities were plotted. An offset distance value was manually calculated for each time point to align each time point to the centre of the pollen tube tip. The combined average of each channel was calculated from a total of 10 images.

### Yeast expression

Fission yeast strain LD328 was used in this study and general methods to transform into fission yeast cells were as previously described (Gietz, 2014). Briefly, the plasmids were linearized with *NotI* restriction enzyme. Yeast cells were cultured by YES medium at 30°C, 200 rpm until OD_600_ to 0.4-0.8, harvested the cells by centrifugation and washed three times with sterile water. Transformation mix (240 µL 50% PEG3350, 36 µL 1.0 M LiAc, 50 µL single-stranded carrier DNA (2.0 mg/mL)) was added to 34 µl linearized plasmid DNA, cells were resuspended by vortexing, and were incubated in a 42°C water bath for 40 min. Transformants were plated on EMM+HT medium (EMM medium supplemented with 45mg/L histidine and 15 µM thiamine). After 3 days, colonies were transferred to EMM+H (EMM medium supplemented with 45mg/L histidine) plates and observed by Zeiss LSM880 confocal laser microscope using a 100× objective.

### *In vitro* protein expression and purification

All proteins were expressed in Escherichia coli BL21 (DE3) cells. Briefly, protein expression was induced by 0.5 mM isopropyl β-D-1-thiogalactopyranoside (IPTG) for 18 h at 18℃. Cells were collected by centrifugation and lysed with buffer A (40 mM Tris-HCl pH7.4, 500 mM NaCl, 10% glycerol). The bacteria were lysed by sonication and the supernatant was flowed through a column packed with Ni-NTA. Proteins were eluted with buffer B (40 mM Tris-HCl pH7.4, 500 mM NaCl, 500 mM Imidazole) and purified with a Superdex 200 increase 10/300 column. Proteins were stored in buffer C (40 mM Tris-HCl pH7.4, 500 mM NaCl, 1 mM DTT) at -80°C.

For the phase separation experiments *in vitro*, MBP-EH1(FL)-GFP, MBP-EH1(ΔIDR3)-GFP and MBP-EH1(ΔCCΔIDR3)-GFP was cleaved with TEV protease for 3 hours to remove MBP tag. Proteins were diluted to the desired concentrations in different concentration of NaCl. Droplets in 384-well plate were observed by Zeiss LSM880 confocal laser microscope using a ×63 objective.

For the co-localization experiments *in vitro*, 5 µM EH1(FL)-GFP after MBP cleaved and 5 µM mCherry-TOL6 were mixed with buffer (40 mM Tris-HCl pH7.4, 100 mM NaCl) and incubated on ice for 30 min. Droplets in 384-well plate were observed as described previously. GFP was excited at 488 nm and detected at 491-535 nm, mCherry was excited at 561 nm and detected at 579-650 nm.

### TurboID proximity labelling

PSB-D *A. thaliana* cell suspension cultures were transformed with the AtEH1-linker-TurboID and experiments were performed as previously described (Arora et al., 2020). The TPLATE-linker-TurboID dataset was described before (Arora et al., 2020). To calculate enrichment scores, the datasets were compared to a large set of in-house TurboID experiments using unrelated baits performed using the same conditions. Proteins identified in at least two experiments, which did not occur in the background list, and showed a high (at least 20-fold) enrichment score (NSAF ratio × −log(P value)) versus the large dataset were considered enriched.

### Analysis of disorder and prion-like regions

For sequence analysis of individual proteins; prion-like residues were obtained using the PLAAC online tool with a core length of 60 and ‘relative weighting of background probabilities’ of 100. Prediction of disordered residues (MobiDB consensus scores) were obtained for the *A. thaliana* proteome (NCBI taxon ID 3702) on 2021-08-31. The *A. thaliana* proteome of prion-like sequences was obtained from previous work (Chakrabortee et al., 2016; Powers et al., 2019). For the classification of endocytic proteins (Figure 4B); proteins were considered disordered if they had a MobiDB consensus score above 0.15 (15% disordered residues), and prion-like if they contained a prion-domain (≥ 60 amino acids) identified in either of the previous published datasets (Chakrabortee et al., 2016; Powers et al., 2019).

### CLEM-ET

The CLEM approach was performed similarly to previously described with minor modifications (Chambaud et al., 2022). For the CLEM-ET of AtEH1 condensates, a small part from the bottom of the hypocotyl was cut from 4-day old *A. thaliana* seedlings stably expressing 35S:AtEH1-GFP. For the rapamycin experiments, seedlings were first treated with 2.5 µM rapamycin for 2 hours in 6 well plates with gentle shaking, and then a small region of the root tip was excised. Root or hypocotyl samples were quickly frozen in a Leica EM-PACT high-pressure freezer using 20% BSA as a cryoprotectant. Freeze-substitution was performed with a Leica AFS 2 in acetone containing 0.1% uranyl acetate. Samples were embedded in HM20 Lowicryl resin. 150nm ultra-thin sections were cut using a Leica Ultracut microtome with a diamond knife (Biel, Switzerland) and placed on copper mesh grids.

EM grids were imaged before ET by fluorescence microscopy using a Zeiss LSM 880 Airyscan confocal microscope with a 63× apochromatic N.A 1.4 oil objective. Transmission electron microscopy observations were carried out on a FEI TECNAI Spirit 120 kV electron microscope equipped with an Eagle 4Kx4K CCD camera. Correlation between fluorescence and electron microscope images was performed using the ec-CLEM plugin in Icy software. For hypocotyl samples, chloroplasts and vacuoles were used as landmarks for registration. For the root samples, cell edges were used as landmarks.

Electron tomography was performed as previously described (Nicolas et al., 2018). Briefly, 5 nm gold beads were briefly applied to grids on both sides and used as fiducials for tomography. Tilt series were acquired from -65° to 65° at 1-degree increments. Pixel size was 0.28 nm or 0.39 nm at 4096×4096 for AtEH1-GFP condensates and rapamycin experiments, respectively. To acquire dual axis tomograms the grid was rotated 90°. The raw tilt series were aligned and reconstructed using the fiducial alignment mode with the eTomo software in IMOD. 10 to 25 fiducials were selected to accurately align images. Reconstruction was performed using the back-projection with SIRT-like filter (10-50 iterations) in IMOD.

### Segmentation and visualisation of EM tomograms

Ribosomes were segmented from tomograms using a pixel and object classification workflow in Ilastik. Ribosomes were manually annotated in 3D using the pixel classification with all available features selected. The resulting probability map was smoothed in 3D (σ=3.5), and a size filter was applied to remove incorrectly annotated ribosomes. The probability map was further refined using object classification with object features selected (excluding location features) to remove incorrectly classified ribosomes. Because of slight differences between contrast between different samples, we performed the segmentation process for individual samples, and applied the resulting classification model to other tomograms using batch processing. Ribosome object predictions were imported into Amira as 8-bit labels together with ET tomography files. Mitochondria and clathrin-coated vesicles were manually segmented in Amira.

### Confocal microscopy

Unless otherwise indicated, all images were acquired using a PerkinElmer Ultraview spinning-disk system, attached to a Nikon Ti inverted microscope, and operated using the Volocity software package. Images were acquired on an ImagEM CCD camera (Hamamatsu C9100-13) using frame-sequential imaging with a 60x water immersion objective (NA = 1.20). Specific excitation and emission were performed using a 488 nm laser combined with a single band pass filter (500-550 nm) for GFP in single camera mode. RFP was visualized using 561 nm laser excitation and a 570-625 nm band pass filter. Exposure time was between 25-500 ms.

For the analysis of AtEH1 truncations or mutations in tobacco the relative saturation concentration was obtained by quantifying the mean cytosolic signal after subtraction of background signal. The partitioning assay quantification was performed using the MitoTally script using regions of interests determined from AtEH1 positive foci (Winkler et al., 2021).

### Whole-mount immunofluorescence imaging

Roots from 4-day-old Arabidopsis seedlings were analysed by immunofluorescence as previously described using an InsituPro Vsi II (Intavis) automated system (Sauer et al., 2006). Antibody dilutions were used as follows: rabbit anti-AtEH1 [1:600], rabbit anti-CHC [1:300], mouse anti-RFP [1:300], Alexa488-conjugated anti-rabbit IgG [1:600], Cy3-conjugated anti-mouse IgG [1:600].

### FM4-64 uptake imaging and analysis

Whole 5-day-old seedlings were incubated with 2 µM FM4-64 (Invitrogen) solution in half-strength MS liquid medium without sucrose at room temperature for 15 min. Samples were briefly washed twice with ½ MS before imaging. Images were acquired using the Leica SP8X system using a ×40 W 1.2 NA objective using 488 nm laser excitation, and collection from 650nm - 750nm. The ratio of plasma membrane signal and cytosolic signal were calculated using an automated detection macro in ImageJ ‘detect_membrane_and_cytoplasm_from_a_roi.ijm’ (Grones et al., 2022), using a threshold set to 60.

### Fluorescence recovery after photobleaching (FRAP)

Tobacco cells transiently expressing proteins were incubated with Latrunculin B (4 μM) for 30 minutes prior to imaging to inhibit condensate movement. FRAP was performed on the spinning disk system using the Ultraview PhotoKinesis unit. Three individual spots were bleached simultaneously, using the ‘stamp’ tool in Volocity to select a square region of 6×6 µm. Bleaching was performed using 100% of the 488 laser for 3s, and post-bleach acquisition was performed every 0.6 seconds for 120 seconds. Fluorescence intensity signal from the condensate was measured by tracking using Trackmate to correct for minor sample drift in x and y axis. Samples which drifted in the z-axis were excluded from analysis. Analysis was performed using the easyFRAP online web tool, using the condensate signal as the ‘region of interest’, an unbleached region as the ‘whole cell area’, and a background region as the ‘background’. The full scale normalisation method was used, and curves were fitted using the double exponential equation.

For the 2in1 ratiometric FRAP experiment, AtEH1 WT and mutant reporters were expressed from a single T-DNA. FRAP image acquisition parameters were the same as above, except images were acquired sequentially at 1 second per two-channel frame. The t_1/2_ times from easyFRAP from GFP and mCherry were divided to get a t_1/2_ ratio (GFP/mCherry). The absolute t_1/2_ values obtained from tobacco experiments are highly variable between samples due to differences in expression levels and cell concentrations. However, the t_1/2_ ratio was relatively consistent despite large differences in absolute t_1/2_ times. To generate visually comparable FRAP curves (Figure 7C), we therefore selected 8 samples from each condition binned to a t_1/2_ time of 17.2 seconds and plotted the resulting curve using easyFRAP.

For the FRAP experiments *in vitro*, MBP-EH1(FL)-GFP was cleaved with TEV protease for 3 hours to remove MBP tag. EH1-FL protein was diluted into 10 µM at 100 mM NaCl. After 30 min of incubation on ice, droplets were bleached with a 488-nm laser pulse (3 repeat, 100% intensity) on Zeiss LSM880 confocal laser microscope using a ×63 objective. Fluorescence recovery was recorded every 2 s for 400 s after bleaching. Images were acquired using ZEN software. Using Fiji/ImageJ to analyse the fluorescence intensity of recovery.

### Condensate tracking

Tracking of condensate fusion (Figure 1D), FRAP (Figure 1E, 7C) and mobility (Figure 3C and S3A) was performed using the Trackmate using the DoG detection algorithm. For the FRAP assay, the ‘extract track stack’ feature in Trackmate was used to generate the individual images.

### Endocytosis imaging and quantification of lifetime

Endocytosis dynamics were imaged on UltraView spinning-disk system using a Nikon Perfect Focus System (PFSIII) for Z-drift compensation. Images of root epidermal cells of 4-day old seedlings expressing pEH1:EH1-GFP were acquired using a 100x oil-immersion objective (Plan Apo, NA = 1.45), with scan settings set at ‘High Speed’. Movies were acquired at 2 frames per second for a duration of 5 minutes.

We excluded cells with very high density, or high background signal as the signal-to-noise ratio was too low for automated quantitative analysis. We imaged epidermal cells within 1-4 cells from the root hair initiation, due to having best S/N ratio here.

For post-processing images in ImageJ were first combined to 1 frame/second using a grouped Z-projection (SUM intensity), cropped to include only the plasma membrane region in focus, contrast adjusted using the ‘auto’ function, then converted to 8bit. Quantitation of images using a modified cmeAnalysis script (Aguet et al., 2013) (Johnson et al., 2021). Tracking parameters (cmeAnalysisTrackingAndDetection) were modified from default parameters as follows to improve tracking accuracy: TrackingGapLength [1], TrackingRadius [1 2].

### Surface rendering of condensates

Time-lapse images of AtEH1 IDR mutant condensates in *N. benthamiana* were acquired with the spinning disk system using a 60x objective using a 1.5x additional zoom with scan settings set at ‘High Speed’. Z-stacks were acquired using 15 slices (0.5 µm spacing) at 2.54 seconds per stack, for a total duration of 4 minutes. Sample drift was corrected in 3D using the ‘Correct 3D drift’ plugin in Fiji. Images were imported into Imaris, and condensates were segmented using the ‘spots’ tool, coloured by ‘Sphericity’. Note that the condensates in *N. benthamiana* have reduced sphericity in 3D due to being pressed between the plasma membrane and the vacuole (which occupies the majority of the cellular volume).

### b-isox enrichment

5-day old Col-0 seedlings were ground in liquid nitrogen, and total proteins extraction and subsequent b-isox enrichment were performed as previously described (Zhang et al., 2022).

### SDS–PAGE and immunoblotting

For pEH1-AtEH1-GFP samples: Samples with 1× Laemmli loading buffer (BioRad) and 1× NuPage reducing agent (Invitrogen) were heated for 10 min at 95 °C. Samples were loaded and separated on a 4–20% SDS–PAGE TGX gel (BioRad) and subsequently blotted on polyvinylidenedifluoride (PVDF; BioRad). Membranes were blocked overnight at 4C in 5% skimmed milk in PBS-T. Next, the blots were incubated with the primary antibodies (α-TPLATE (Yperman et al., 2021b), 1:1000], α-AtEH1/Pan1 (Grones et al., 2022), 1:1000], α-CHC [Agrisera, AS10690, 1:2000], α-Tubulin [Sigma-Aldrich, T5168, 1:5000] and secondary antibodies (α-rabbit-HRP [Cytiva, NA934, 1:10000], α-mouse-HRP [Cytiva, NA931, 1:10000]) in 3% skimmed milk in PBST for 1 h. HRP-conjugated antibodies were detected using western lighting plus-ECL reagent (Perkin elmer).

### Multiple sequence alignment and phylogenetic analysis

To identify AtEH1/Pan1 homologues, predicted proteins of selected genomes from the Phytozome v13 database were searched using the BLASTP algorithm with Arabidopsis AtEH1 as an input sequence. Multiple sequence alignment was constructed with the MAFFT algorithm in the einsi mode. Phylogenetic analysis was carried out utilizing PhyML v3.0 with the smart model selection. The phylogenetic tree was visualized using iTOL v6.

### Statistical analysis

Statistical analysis was performed using Graphpad Prism. Data was tested for normality for analysis. Significance criterion was set at a p value of <0.05.

### Data availability

All materials are available from the corresponding authors upon request.

## Dragwidge *et al.,* (2023): Biomolecular condensation orchestrates clathrin-mediated endocytosis in plants

### Supplemental information

Figure S1 to S7

Video S1 (Related to Figure 3)

Video S2 (Related to Figure 5)

Video S3 (Related to Figure 6)

Video S4 (Related to Figure 6)

Video S5 (Related to Figure 7)

Table S1. Detailed list of cloning vectors

Table S2. Mass spectrometry data

**Figure S1 (Related to Figure 1).**
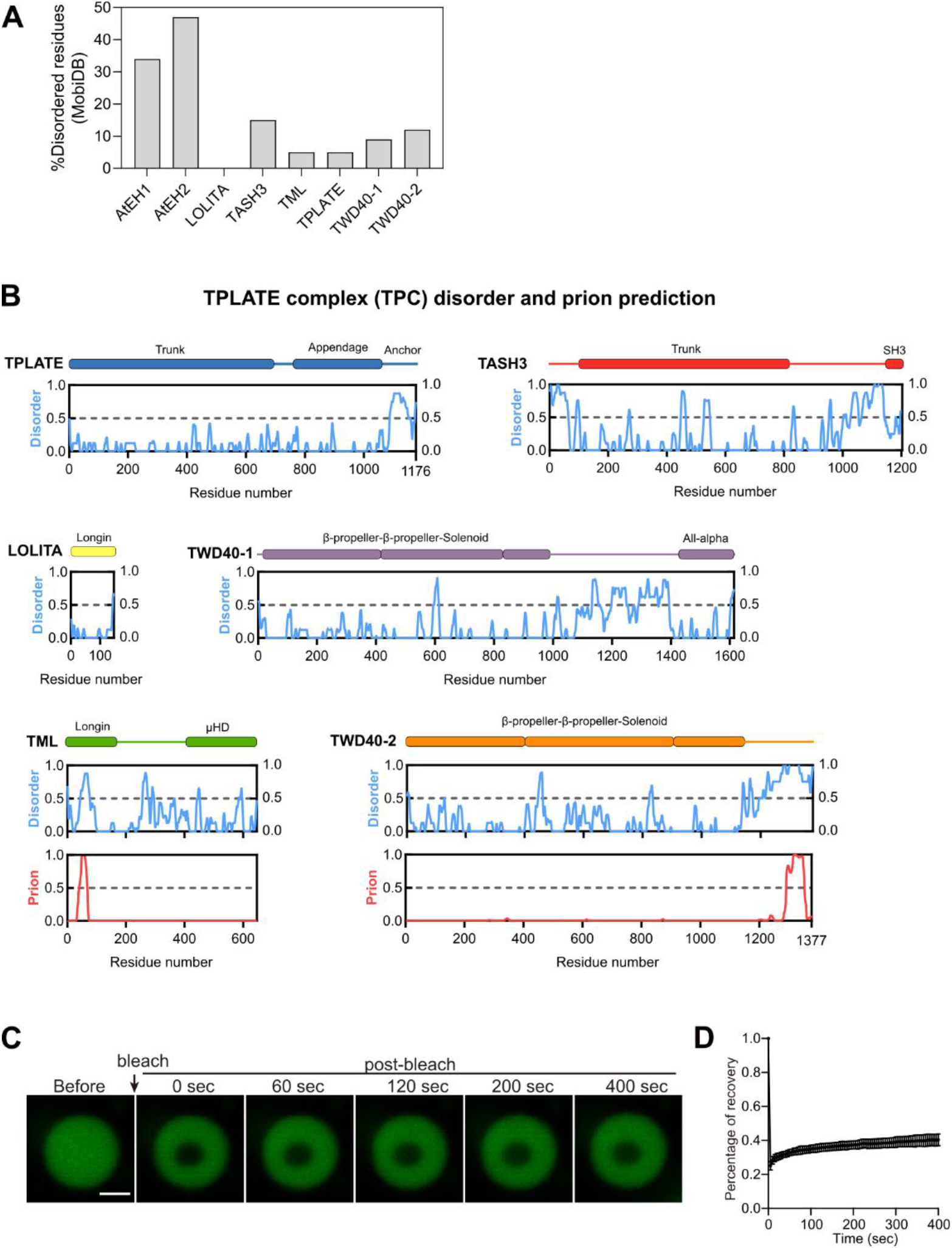
TPC disorder prediction, and AtEH1 *in vitro* FRAP assay. (A) Plot of the proportion of disordered residues for TPC subunits. (B) Plot of TPC subunits showing prediction of disordered (MobiDB, blue), and prion-like (PLAAC, red) residues. Regions with values >0.5 are considered disordered, or prion-like respectively. Prion-like residues were only identified in TML and TWD40-2 subunits. (C-D) *In vitro* FRAP assay of purified GFP-AtEH1_FL_ protein. Scale bar = 2 μm.

**Figure S2 (Related to Figure 2).**
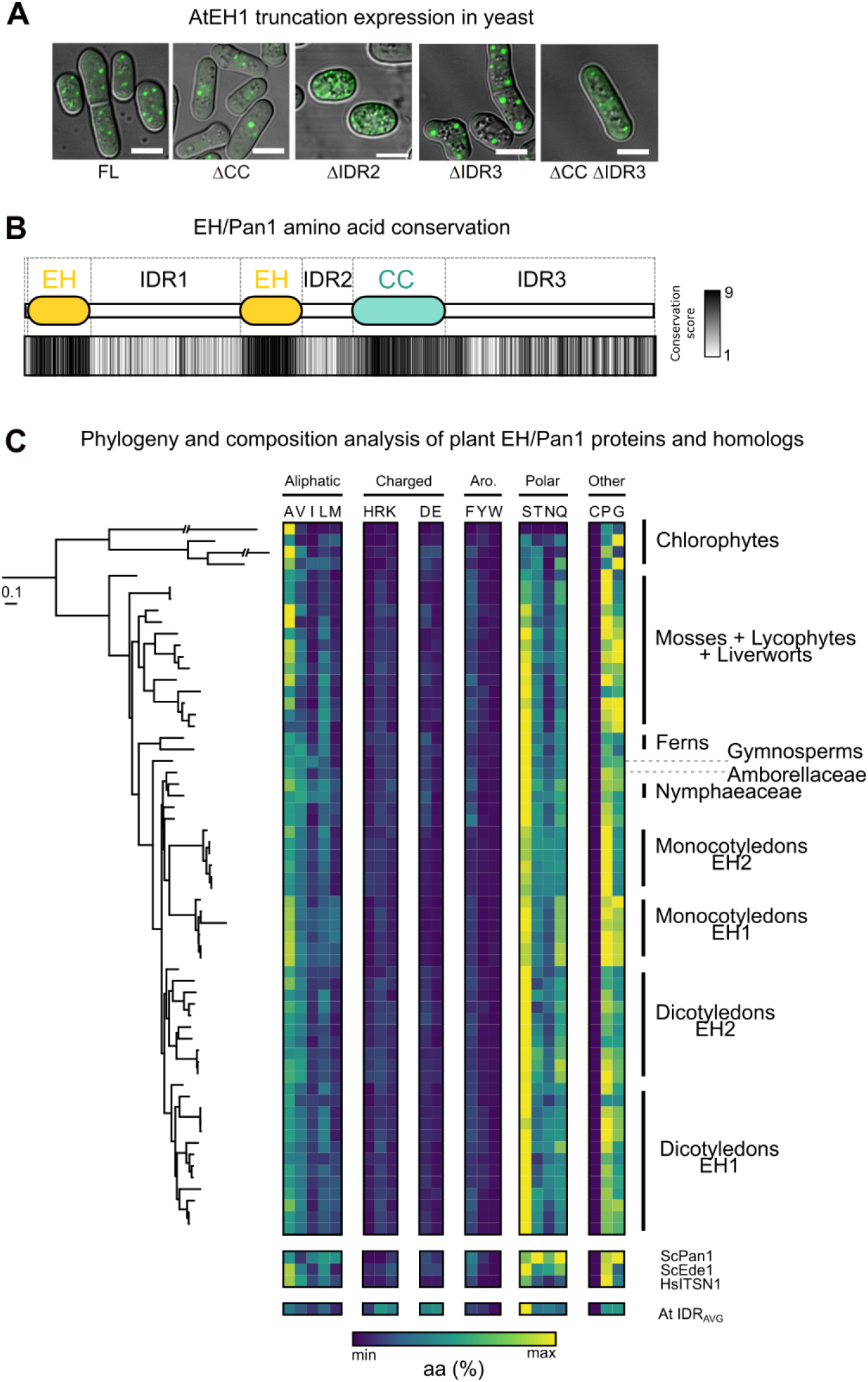
AtEH1 truncation construct expression in yeast, and evolutionary comparison of EH/Pan1 IDR1 amino acid composition across Archaeplastida. A) Expression and condensate formation capacities of AtEH1 domain constructs in *S. pombe*. Scale bars = 5 μm. B) Conservation of amino acids at the single residue level based on Consurf analysis calculated using 128 AtEH homologous sequences throughout plant evolution. The average conservation score is indicated for each region (1 = 0% conservation, 9 = 100% conservation). IDR1 is highly variable at the individual amino acid level. C) Phylogenetic tree representing the maximum likelihood phylogeny of selected EH proteins. The phylogenetic tree was arbitrarily rooted to reflect phylogenetic relationships between chlorophyte and streptophyte lineages. The scale bar represents 0.1 amino acid substitution per site. The composition of the IDR1 for EH/Pan1 proteins is shown, with each amino acid plotted with the total number of each residue indicated. EH/Pan1 homologs from yeast (ScPan1, ScEde1), human (ITSN1), and the *A. thaliana* proteome average IDR are also indicated. The sequences used to construct the alignment in panel B and the phylogenetic tree in panel C can be found in the source data.

**Figure S3 (Related to Figure 3).**
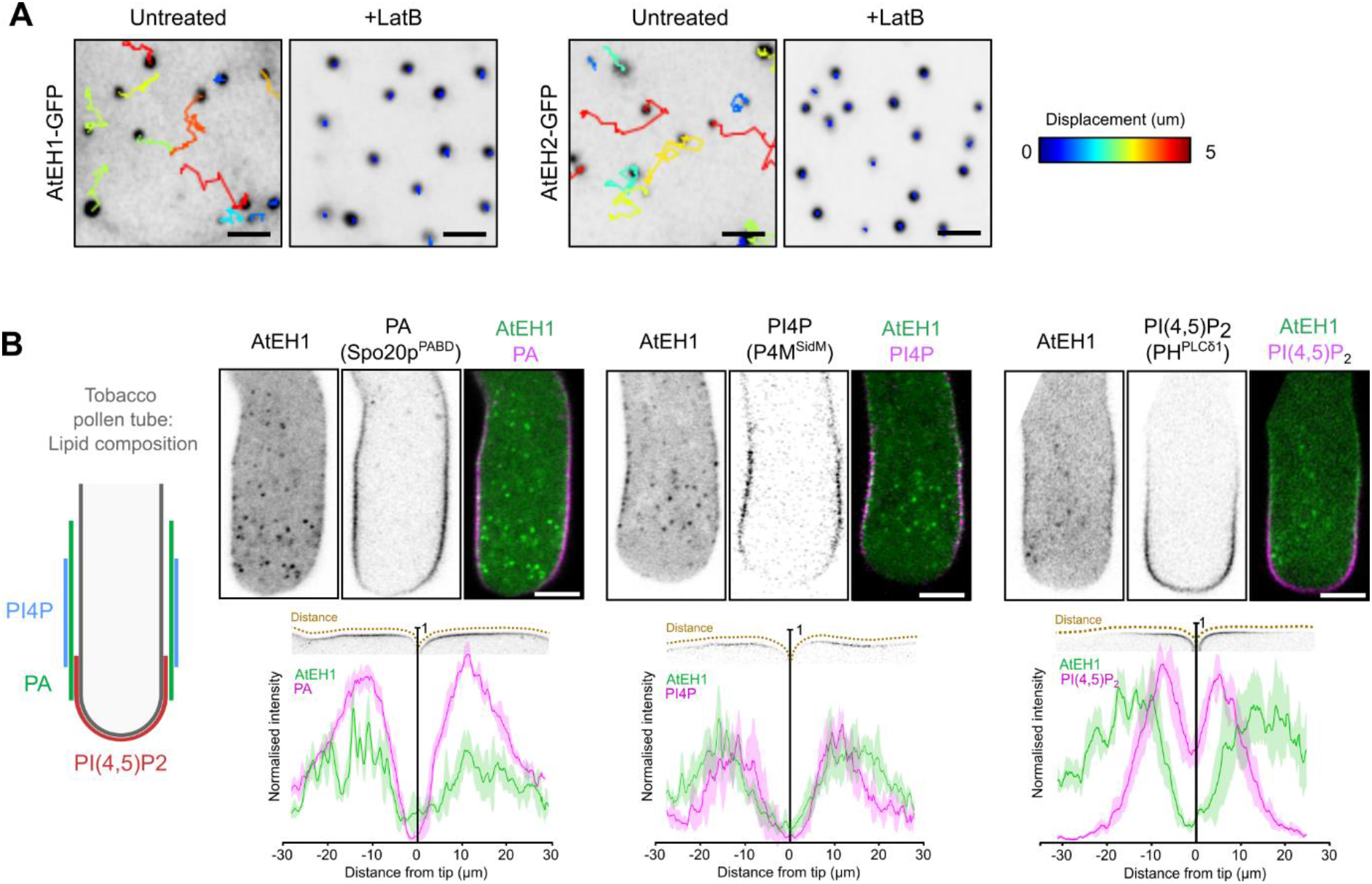
Condensate motility and phospholipid specificity experiments. (A) Time-lapse and tracking of AtEH1 and AtEH2 condensates in Latrunculin B treated (4 μM, 30 minutes) *N. benthamiana* epidermal cells. (B) Schematic representation of the phospholipid localisation domains in Tobacco pollen tubes. Images show transient co-expression of pLAT52:AtEH1-YFP with lipid biosensors (pLAT52-biosensor-mCherry) in *N. tabacum* pollen tubes. The chart shows the quantification of the plasma membrane signal from AtEH1 and lipid biosensors from the tip of the pollen tube. Data represent mean ± SD from five time points from two individual pollen tubes. Scale bars = 5μm.

**Figure S4 (Related to Figure 4).**
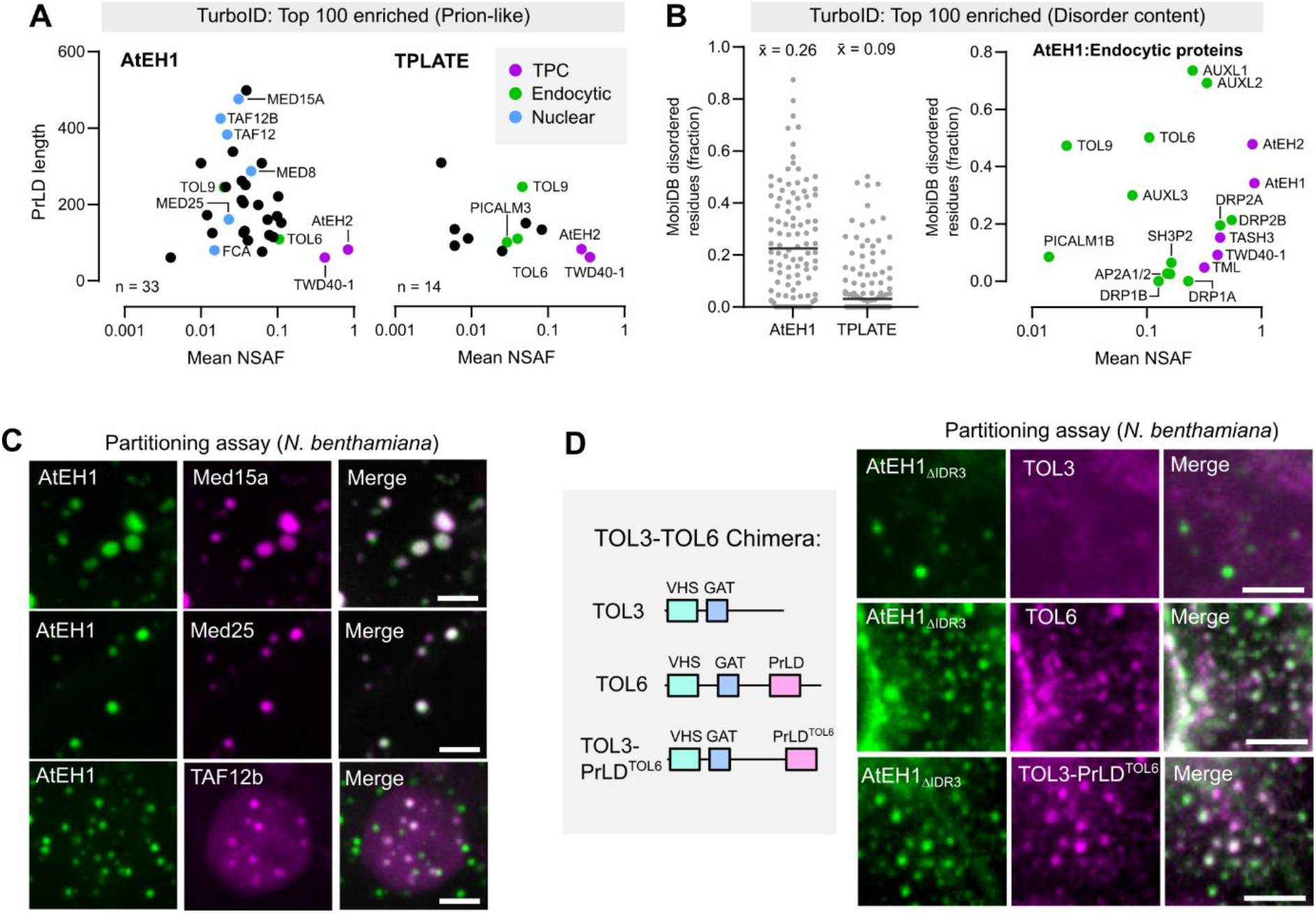
AtEH1 interacts with and recruits proteins with prion-like domains. (A-B) Plot of Prion-like proteins identified in the AtEH1 and TPLATE TurboID datasets (A). Plot of proteins based on disorder content (MobiDB) and plot showing abundance and disorder content of selected endocytic proteins (B). 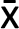 (mean) disorder score is indicated. (C) Partitioning assay of selected nuclear proteins with prion-like domains identified in the AtEH1-TurboID dataset. UBQ10:AtEH1_FL_-GFP was co-expressed with mScarlet tagged client proteins in *N. benthamiana* epidermal cells. Partitioning was observed in the cytosol (Med15a, Med25) and in the nucleus (TAF12b). (D) TOL chimera experiment. The Prion-like domain (PrLD) of TOL6 was added onto TOL3 to generate TOL3-PrLD^TOL6^-mScarlet. The different constructs were co-expressed with AtEH1_ΔIDR3_ in *N. benthamiana* and imaged. The prion-like domain of TOL6 is sufficient to partition TOL3 with AtEH1_ΔIDR3_. Scale bars = 5 μm (C, D).

**Figure S5 (Related to Figure 5).**
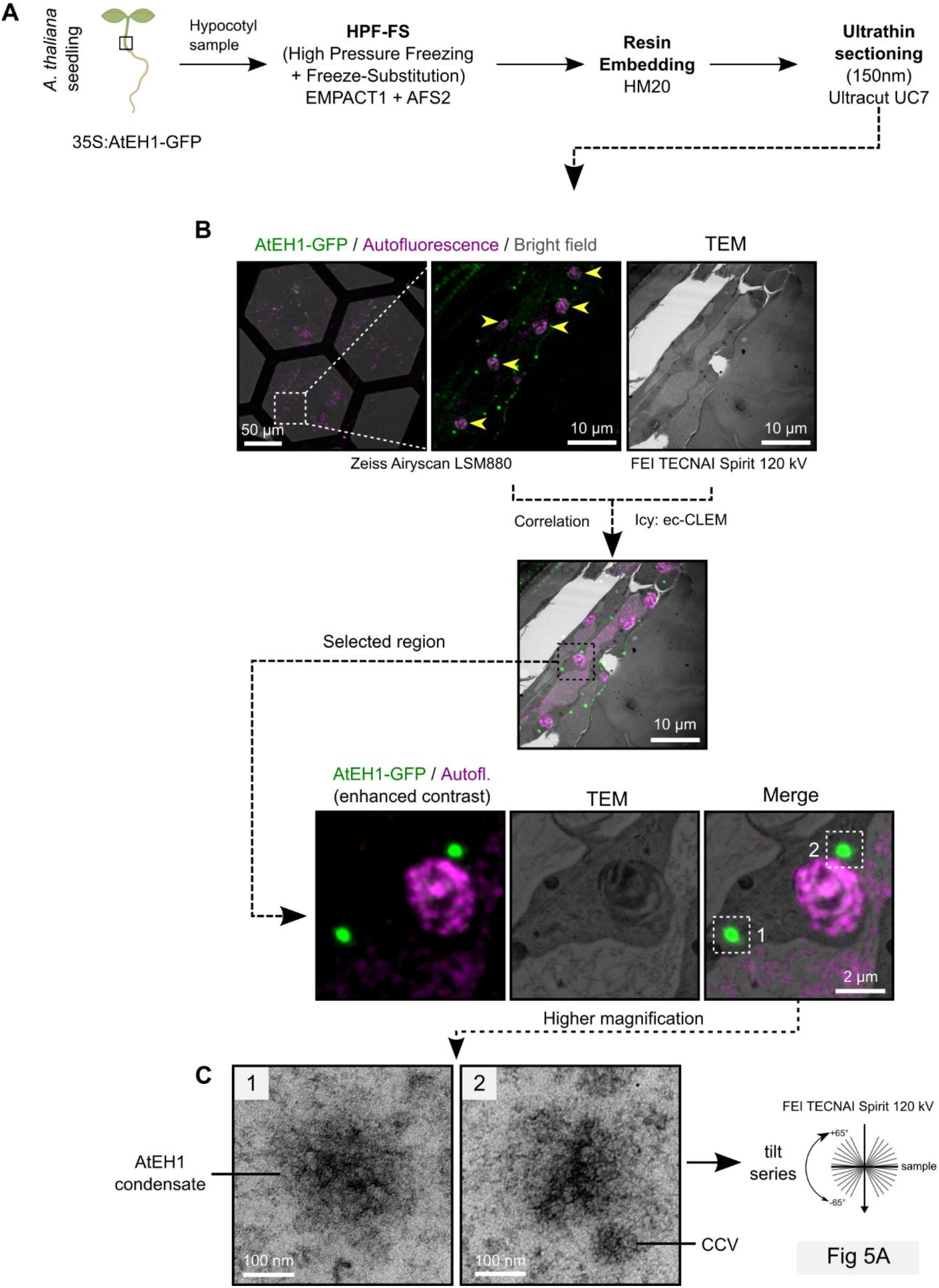
CLEM-ET workflow. (A) Overview of the sample preparation procedure. (B) Correlative light and electron microscopy imaging of hypocotyl sections. Yellow arrowheads indicate chloroplasts used as natural landmarks for correlation. (C) Zoom in of regions selected for ET reconstruction. The condensates have similar electron density to CCVs.

**Figure S6 (Related to Figure 6).**
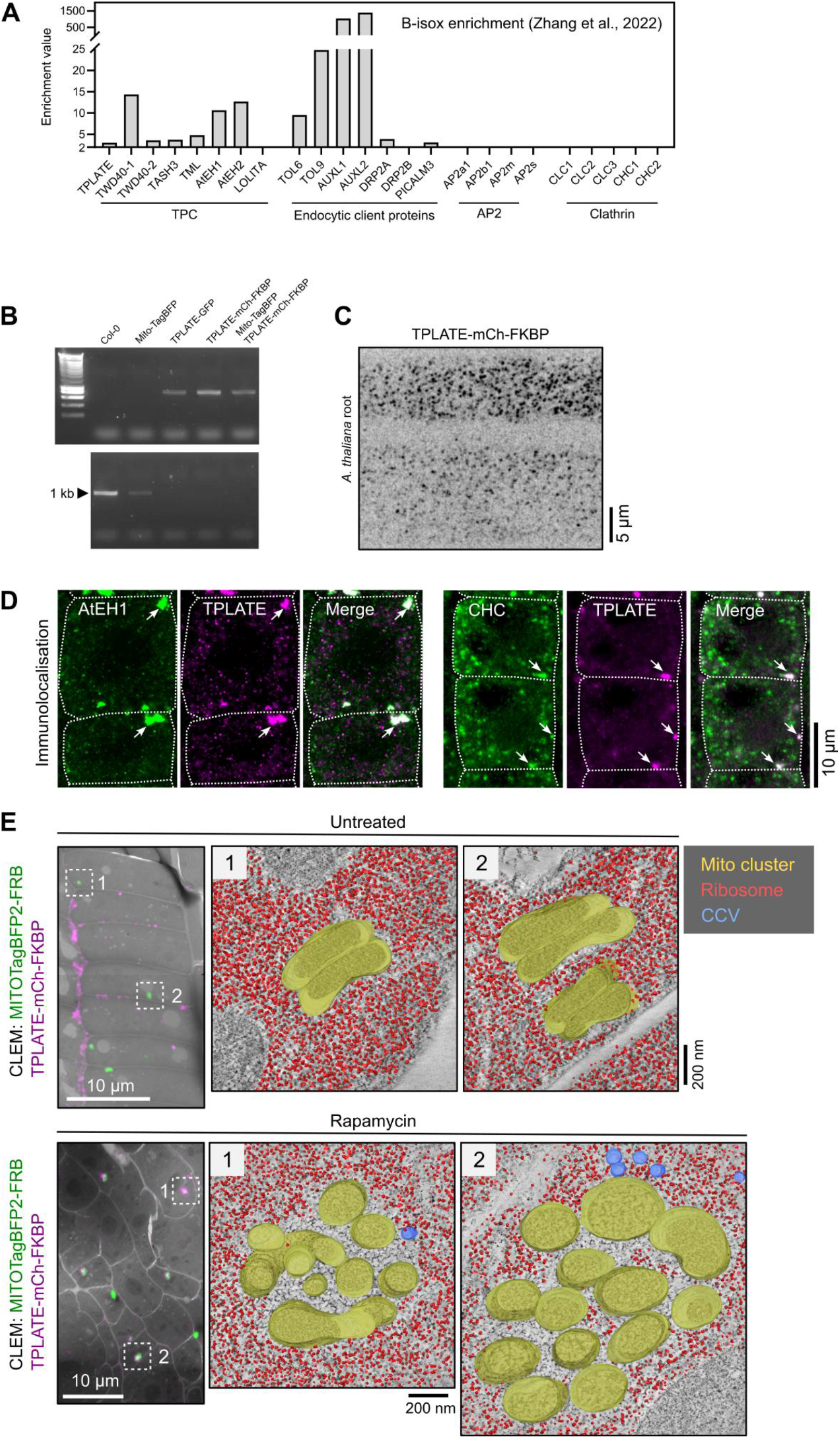
B-isox enrichment and TPLATE re-localisation additional data. (A) Enrichment of TPC subunits and endocytic proteins after b-isox treatment. Data shown is from Zhang et al., 2022. Proteins with enrichment values ≥ 2 are considered enriched, and < 2 not enriched. Enrichment values below 2 were not represented in Zhang et al., 2022. (B-C) Validation of TPLATE-mCh-FKBP *tplate A. thaliana* lines. Genotyping PCR indicate TPLATE-mCh-FKBP complements the *tplate* mutant (B). Top row: *tplate T-DNA*, bottom row: WT TPLATE; TPLATE-GFP *tplate*(-/-) (positive control). Endocytic foci from *A. thaliana* root cells expressing TPLATE-mCh-FKBP (C). (D) Whole mount immunolocalisation of AtEH1 (Anti-AtEH1), CHC (Anti-CHC), and TPLATE-FKBP-mCherry (Anti-RFP) in rapamycin treated *A. thaliana* root cells. Arrows indicate mitochondria clusters. (E) CLEM localisation of untreated and rapamycin treated mitochondria clusters (left), and ET reconstruction representing the full segmented 3D volume. Ribosomes (red) are excluded from the interior of the mitochondria clusters (yellow) in rapamycin treated root cells. No exclusion zone was observed in untreated cells. CCVs (blue) are observed nearby mitochondria clusters in rapamycin treated root cells.

**Figure S7 (Related to Figure 7).**
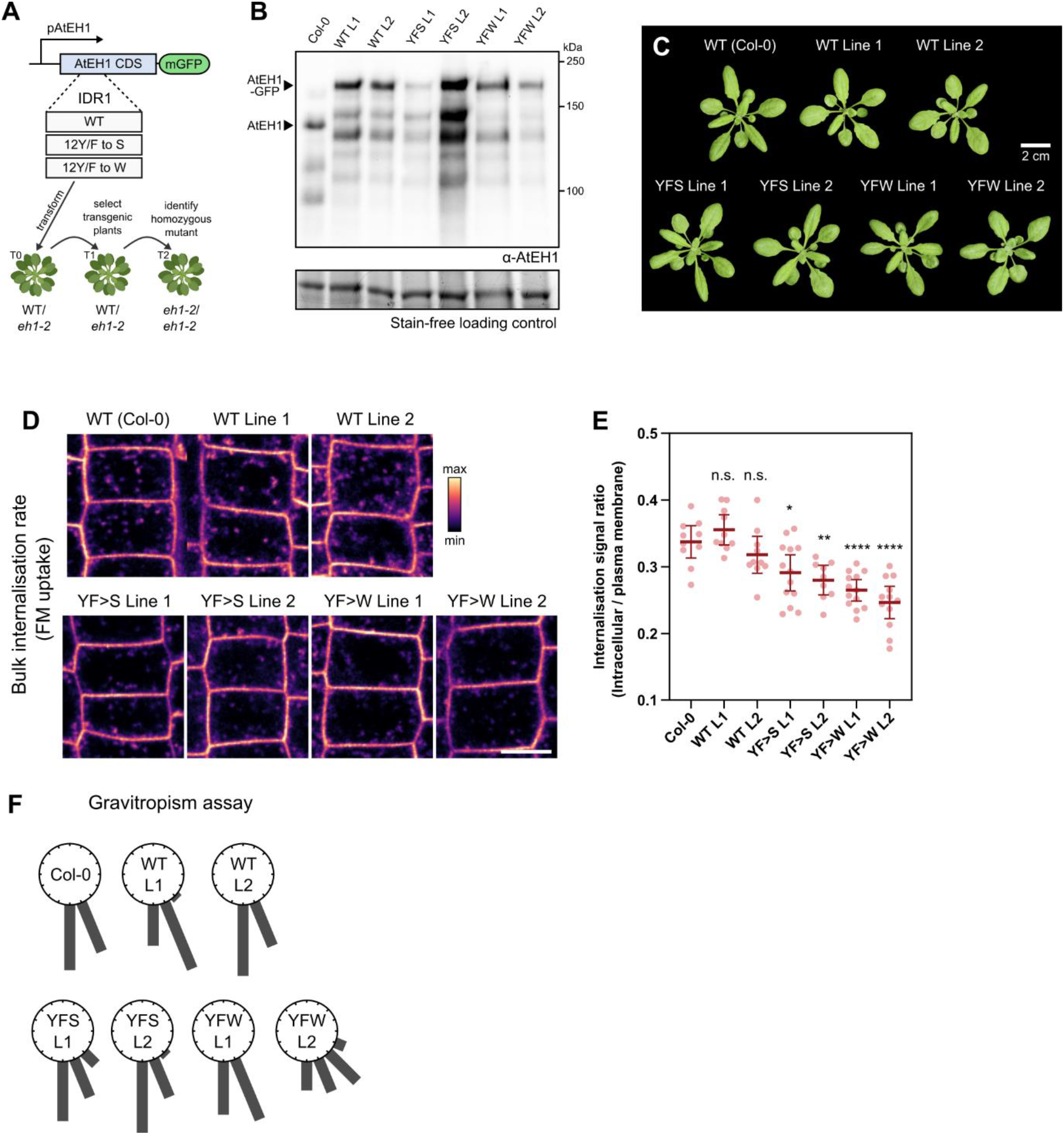
Generation and validation of pAtEH1:AtEH1-GFP reporters. (A) Schematic of the generation of pAtEH1:AtEH1-GFP rescue lines. (B) Western blot of *A. thaliana* protein extracts using an antibody directed against AtEH1, indicating absence of endogenous AtEH1 in the recued lines. Lower molecular weight bands are degradation products. Strain free gel is shown as a loading control. (C) Representative plant rosette image of WT Col-0 and AtEH1 rescued lines. (D-E) Endocytic flux experiment. Seedlings were treated with the styryl dye FM4-64 (2µM, 10min) and root cells were imaged. (E) Quantification of FM4-64 internalisation by measuring the ratio of intracellular versus plasma membrane intensity. Bars indicate mean ± 95% CI. Statistics indicate significance to WT; n.s. not significant, * p<0.05, ** p<0.01, **** p<0.0001, unpaired t-test. (F) Gravitropism assay. Roots grown on ½ MS agar plates were gravi-stimulated by turning them 90°. The gravitropism plot indicates the angle of the root tip in 22.5° bins. n= 20 roots for each genotype. Scale bar = 10 μm (D).

